# Exogenous application of KNO_3_ elevates the salinity tolerance of *Stevia rebaudiana* through ion homeostasis mechanism

**DOI:** 10.1101/657767

**Authors:** Mitali Mahajan, Surbhi Sharma, Pawan Kumar, Probir Kumar Pal

## Abstract

Though relatively little is understood of adaptation, physiological and metabolic changes of *Stevia rebaudiana* under exposure to salinity stress, it is hypothesized that exogenous application of potassium (K^+^) could elevates the salinity tolerance through ions homeostasis. Thus, an experiment was conducted with twenty treatment combinations comprising four salinity levels (irrigation with normal water as control and three level of NaCl at 40, 80 and 120 mM) and five different concentrations of KNO_3_ (0.0, 2.5, 5.0, 7.5, and 10.0 g L^−1^). Dry leaf yield was not negatively affected with mild salinity (40 mM). However, the detrimental effects were observed at moderate and higher salinity levels (80 and 120 mM). The uptakes of K^+^, Ca^2+^, and N were significantly reduced at higher salinity level, whereas accumulations of Na^+^ and Cl^−^ ions in plant tissues were substantially increased. Proline content in leaf was also increased significantly (*P*≤0.05) in response to salt stress. Among the foliar application, KNO_3_ at 5.0 gL^−1^ registered significantly (*P*≤0.05) higher dry leaf yield compared with control. Exogenous application of K^+^ under moderate salinity stress maintained ion balance in cytosol, particularly K: Na. Thus, the salinity tolerance of stevia can be elevated to some extent through exogenous application of K^+^.

**Highlight:** The detrimental effects of moderate and higher salinity levels on growth and dry leaf yield of stevia were observed. However, tolerance level can be elevated through exogenous application of KNO_3_.

## 1. Introduction

The burgeoning rate of diabetic and obesity patients is a serious apprehension in the worldwide. About 346 million people worldwide are in the grip of diabetes whereas in India 69.2 million people are diabetic (WHO 2015). The cane sugar is not recommended for consumption to the diabetic and obesity patients. In these circumstances, stevia (*Stevia rebaudiana*) has emerged to be a natural sweet gift for millions of diabetics as a sweet quotient in their daily life. The stevia leaves contain sweet-tasting and low-calorie diterpenoid steviol glycosides (SGs), which are nearly 300 times sweeter than sucrose (Kinghorn 2002). However, the performance of stevia in saline soil or plant irrigated with saline water is not properly elucidated so far. The global extent of salt-affected land amounts to about 1128 million ha (FAO 2008). The term salt affected refers to the soils that are saline or sodic, which is one of the major stress factors for agriculture in consequence of climate change. Salinity stress alone adversely affects to the production in over 30% of irrigated crops and 7% of dryland agriculture worldwide (Schroeder *et al.* 2013). Development of agrotechniques to elevate the salt-tolerance within plant or cultivation of salt-tolerant crops may be an efficient approach for better utilization of salt-affected land to address the related issues.

The restriction of plant growth in the saline soil is directly linked to osmotic potential of the soil solution and total concentration of soluble salts, and the different mechanisms have been developed by the plants to cope with these situations (Munns 2002; Tavakkoli *et al.* 2011). The presence of higher concentration of NaCl in soils increases the accumulation of the phytotoxic ions sodium (Na^+^) and chloride (Cl^−^) in plant organs, concurrently reduces the uptake of other essential nutrients, such as potassium (K^+^), calcium (Ca^2+^), magnesium (Mg^2+^) and nitrate (NO_3_^−^) ion (Niu *et al.* 1995; Cornillon and Palloix, 1997; Halperin *et al.* 2003; Munns and Tester 2008; James *et al.* 2011). The imbalance of ions creates the decisive conditions for plant survival by intercepting different plant mechanisms. Thus, plant accumulates various osmolytes (proline, glycine betaine and sugars), secondary metabolites and antioxidants in response to stress which helps in the survival of the plant through maintaining plant turgor (Ashraf and Foolad 2007).

Salt stress restricts the N assimilate in plant body through the attenuation of activity of cytosolic NADH nitrate reductase enzyme in nitrate assimilation pathway (Sivasankar and Oaks 1996; Jabeen and Ahmad 2011). Potassium is another important macronutrient for optimum growth and development of the plant. Potassium also plays important role in maintenance of osmotic adjustment during stress conditions. It is established fact that K^+^ concentration in plant tissue declines as the Na^+^ concentration in the root media is increased (Subbarao *et al.* 1990; Izzo *et al.* 1991; Perez-Alfocea *et al.* 1996). This disruption of K^+^ further disturbs many metabolic processes in the cytoplasm (Marschner 1995).

Thus, the mechanisms to salinity tolerance should be understood properly to elevate salt tolerance for a particular plant species for cultivation in salt affected areas. Exclusion of surplus Na^+^ or compartmentalization and ion homeostasis are the basic mechanisms of salt tolerance for plants (Li *et al.* 2006; Munns and Tester 2008). So, the exogenous application of nutrients particularly K^+^ and NO_3_^−^ through foliar feeding under saline conditions may be an effective mechanism to reduce Na^+^ and Cl^−^ injury and to elevate tolerance against saline to various extents. Foliar spray is an efficient approach, which involves active and passive processes for translocation of nutrients through leaves to other organs of plant when nutrient uptake from soil is disturbed (Fageria *et al.* 2009).

However, the degree of salinity tolerance depends upon plant spices. Very few research works have been conducted to understand the effect of salinity on growth, the extent of tolerance and mechanism of tolerance in stevia. Similarly, the attempt toward elevating the salinity tolerance of stevia has not been initiated so far. Thus, the present experiment was executed with the specific objectives of (i) determining the degree of tolerance and understanding the effects of salinity on the growth, biochemical activities, and steviol glycoside accumulation; (ii) evaluating the effectiveness of exogenous application of K and N to elevate the salinity tolerance in *S. rebaudiana.*

## 2. Materials and Methods

### 2.1. Experimental location, plant material and application of treatments

A pot experiment was conducted in the open-ended poly-tunnel at CSIR-Institute of Himalayan Bioresource Technology, Palampur, India during two growing seasons of 2016 and 2017. The physico-chemical properties of the experimental soil are presented in **Table 1**. Sixty-days-old stevia seedlings were selected randomly from uniform populations maintained in a poly-house under natural light for transplantation into the pot containing 14 kg soil. For basal dose, 690 mg N as urea, 136 mg P as single super phosphate, and 311 mg K were applied to the pots. At twentieth-days of transplantation, plants were taped in order to maintain the uniform height. The experiment was carried out in two-factor factorial complete randomized design (CRD) with ten replications. The study includes twenty treatment combinations involving four salinity levels of irrigation water (irrigation with normal water as control and three level of NaCl at 40, 80 and 120 mM) and five different concentrations of KNO_3_ (water spray as control, KNO_3_ at 2.5, 5.0, 7.5, and 10.0 g L^−1^). After transplanting, plants were irrigated with normal tap water. Thereafter, individual plants were irrigated treatment wise as per requirement throughout the experimental period. The foliar spray of KNO_3_ for different treatments was initiated at 10^th^ day after first salinity treatment. Distilled water was used for water spray as control treatment. The foliar spray of KNO_3_ solutions took place at the interval of 10 days for three times.

**Table 1.** Physico-chemical properties of the experimental soil

| Parameters | Year |  |
| --- | --- | --- |
|  | 2016 | 2017 |
| Sand (%) | 42.4 | 42.7 |
| Silt (%) | 42.7 | 38.0 |
| Clay (%) | 17.3 | 19.3 |
| Texture | Loam | Loam |
| Bulk density ($\text{g cm}^{-3}$) | 1.25 | 1.17 |
| pH (1:2) | 6.72 | 6.31 |
| EC<sub>25</sub> (1:2)($\text{ds.m}^{-1}$) | 0.11 | 0.10 |
| OC (%) | 1.53 | 1.66 |
| N ($\text{mg kg}^{-1}$) | 64.87 | 122.73 |
| P ($\text{mg kg}^{-1}$) | 14.96 | 13.21 |
| K<sup>+</sup> ($\text{mg kg}^{-1}$) | 272.80 | 284.30 |
| Ca<sup>2+</sup> ($\text{mg kg}^{-1}$) | 925.00 | 939.80 |
| Na<sup>+</sup> ($\text{mg kg}^{-1}$) | 36.80 | 42.00 |
| Cl<sup>-</sup> ($\text{mg kg}^{-1}$) | 3.54 | 3.12 |

### 2.2. Growth and yield

The five plants were randomly harvested after 90 days of transplantation from each treatment. After recording plant height (cm) and number of branches per plant, the leaves were separated from the stems. Leaf area (LA) of individual plant sample was measured with the help of Windias-3 Image analysis system (Delta-T Devices Ltd. UK). The fresh weight of leaf, stem and root were also recorded immediately. Then, the samples were dried at 70 °C± 2 °C in hot air oven until constant weight was attained in order to measure the dry weight. Specific leaf weight (SLW) was also calculated based on leaf area and dry leaf weight, and it was expressed in mg cm^−2^. The dried leaf, stem and root samples were stored for further chemical and biochemical analysis.

### 2.3. Chlorophyll (Chl) analysis

The leaves were collected from each pot at harvesting stage for estimation of Chl content in the leaf. The Chl was extracted from 200 mg fresh leaf samples with a 25 mL of 80% acetone (v/v) and was kept under dark conditions at room temperature for 24 h to avoid photooxidation. The absorbance of the extract was recorded at two different wavelengths i.e. 645 nm and 663 nm with a UV/Vis spectrophotometer (Shimadzu, UV-2600). The Chl_*a*_, Chl_*b*_ and total Chl content (mg g^−1^) were determined by using standard equations given by Arnon (1949).

### 2.4. Determination of Proline

The accumulation of proline in leaf was quantified as per the method recommended by Bates *et al.* (1973). The proline was extracted from 0.5 g fresh leaf samples of stevia with 10 mL of 3% sulphosalicylic acid and then, centrifuged at 12000 g for 10 minutes. 1 mL supernatent was reacted with the equal volume of acid-ninhydrin and glacial acetic acid in a test tube and incubated for 1 h at 100° C. The reaction was stopped by keeping the test tube in an icebath. Then, 2 mL tolune was added to test tube and mixed vigorously and left it undisturbed for 30 minutes at room temperature. After that, the sample mixture was separated into two phases. The optical density of the chromophore containing tolune was measured at 520 nm with spectrophotometer (model T 90 C UV/vis, PG Instrument Ltd.). The proline content was determined based on standard curves developed with D-proline.

### 2.5. Total phenols

For the analysis of total phenols, 0.1 g of leaf sample was suspended in 25 mL solution of 70% acetone in a conical flask. The conical flask was kept on shaking water bath at 37° C for 2 h. After that, the homogenate was centrifuged at 10000 g for 20 minutes. The supernatant was filtered and collected in a test tube. 1 mL of the extract was taken into a 25 mL volumetric flask, then, 0.5 mL of 1 N Folin-Ciocalteu and 2.5 mL of 35% sodium carbonate were added. The final volume was made up to 25 mL with distilled water. After vortex, the flasks were left in dark conditions at room temperature for 40 minutes for colour development. The intensity of purplish-blue colour was measured at 730 nm with spectrophotometer. A standard curve, prepared with gallic acid, was used for the estimation of total phenols content, and results were expressed as mg gallic acid equivalent (GAE) per g dry leaf.

### 2.6. Steviol glycosides analysis

For the estimation of SGs content in stevia leaves, the collected leaf samples were dried at 40± 2° C in a hot air oven until constant weight was achieved. The dried leaf samples were powdered by a grinder. 100 mg of prepared samples were immersed in 10 mL methanol for 24 h, then filtered. The filtrate was vacuum dried under reduced pressure, after that defatted with n-hexane. Defatted sample was dissolved in 10 mL high-performance liquid chromatography (HPLC) – grade acetonitrile and water (1:1) mobile phase, then filtered with micro filter (0.22 μm pore diameter). The filtrate was used for the determination of important eight steviol glycosides i.e. stevioside, Reb-A, Reb-F, Reb-C, dulcoside-A, rubusoside, Reb-B and steviolbioside (SB) by Ultra HPLC system (LC-MS Shimadzu, 2020 system). The system is equipped with LC-30 AD auto detector, quaternary pump, SIL-30 AC autosampler, Degaser Unit DGU-20 A5R, reverse phase C18 column and SPD-M 30A diode array detector. The temperature of column was set at 30° C. The used mobile phase was the combination of A (5 mM ammonium acetate and 12 ppm equivalent v/v of formic acid) and B (pure acetonitrile) with a ratio of 70: 30 and flowed at the rate of 0.24 mL min^−1^ for 20 minutes. The detection wavelength was 210 nm. The steviol glycosides were quantified based on the standard curves developed with the standard sample of stevioside, Reb-A, Reb-F, Reb-C, dulcoside-A, rubusoside, Reb-B, and steviolbioside.

### 2.7. Determination of plant mineral ion in different plant organs

For understanding spatial distribution of N, P, K^+^, calcium (Ca^2+^), Na^+^ and Cl^−^ in different plant parts, the dried leaf, stem and root samples were grounded in a grinder to pass through a 40 mesh screens. The total N content was estimated by micro-kjeldahl method after the digestion of plant samples in concentrated sulphuric acid. For the estimation of other ions (P, K^+^, Ca^2+^, and Na^+^), the samples were digested with a mixture of concentrated nitric acid, sulphuric acid and perchloric acid (9:4:1). The total P in leaf, stem and root samples was determined with a spectrophotometer (model T 90+ UV/ vis, PG Instrument Ltd.), whereas a flame photometer (model BWB XP, BWB technologies UK Ltd., UK) was used for the estimation of total K^+^, Na^+^, and Ca^2+^ as per the protocols recommended by Prasad *et al*. (2006). The Cl^−^ ion was analyzed by the titration method prescribed by Husband and Godden (1927).

### 2.8. Statistical analysis

The growth, yield and biochemical data obtained from this investigation for consecutive 2 years were subjected to analysis of variance (ANOVA) with the help of Statistica 7 software (Stat Soft Inc., Tulsa, Oklahoma, USA) for estimating the variance components of main effects (salinity level and foliar application of KNO_3_) and their reciprocal interactions effects. The least significant difference value was used to separate the treatment means when *F-test* was significant (*P* ≤ 0.05). A second-degree-polynomial regression model was used to establish the relations between salinity level and dry leaf yield, and foliar application of KNO_3_ and dry leaf yield of stevia. Agronomic traits, SGs profile, biochemical traits and ions accumulation were also subjected to the principal component analysis (PCA) to understand those, which were largely influenced by the treatment combinations.

## 3. Results

### 3.1. Yield attributes

The major yield attributes of the stevia viz. plant height, number of branches and LA were significantly (*P* ≤ 0.05) influenced by the salinity levels and foliar spray of KNO_3_ during both the cropping seasons (**Table 2**). Averaged across the KNO_3_ levels, plants treated with high concentration of NaCl (≥ 80 mM) showed significantly (*P* ≤ 0.05) lower plant height, number of branches and leaf area as compared with the plants irrigated with the non-saline water as control. Compared with control, irrigation of high saline water (NaCl at 120 mM) reduced plant height by 10.86 and 12.58 %, number of branches by 24.26 and 31.29 % and LA by 20.90 and 20.89% in 1^st^ and 2^nd^ cropping seasons respectively. On the same time, application of KNO_3_ at 5.0 g L^−1^ registered maximum height (70.18 and 67.58 cm), number of branches (7.17 and 7.33 No. plant^−1^) and LA (390.92 and 407.66 cm^2^), which are significantly (*P* ≤ 0.05) different from the water spray as control in both years. The interaction effects between salinity levels and KNO_3_ on plant height and number of branch were insignificant (*P* ≥ 0.05), but significant (*P* ≤ 0.05) effect was observed in LA during both cropping years (**Table 2**).

**Table 2.** Effect of salinity and foliar application of KNO_3_ on yield attributes of *Stevia rebaudiana*

| Salinity level (NaCl) | Concentration of KNO<sub>3</sub> | Plant height (cm) |  | Total branch (No. plant<sup>-1</sup>) |  | Leaf area (cm<sup>2</sup> plant<sup>-1</sup>) |  | Specific leaf weight (mg cm<sup>-2</sup>) |  |
| --- | --- | --- | --- | --- | --- | --- | --- | --- | --- |
|  |  | 2016 | 2017 | 2016 | 2017 | 2016 | 2017 | 2016 | 2017 |
| Control (S<sub>0</sub>) | Control (K<sub>0</sub>) | 66.00 | 63.33 | 6.67 | 7.33 | 355.36 | 350.64 | 15.42 | 15.20 |
|  | 2.5 g L<sup>-1</sup> (K<sub>1</sub>) | 74.13 | 69.33 | 7.33 | 8.33 | 396.56 | 393.51 | 15.31 | 15.21 |
|  | 5.0 g L<sup>-1</sup> (K<sub>2</sub>) | 74.30 | 71.67 | 7.67 | 8.33 | 460.97 | 467.04 | 15.26 | 15.36 |
|  | 7.5 g L<sup>-1</sup> (K<sub>3</sub>) | 66.57 | 65.00 | 7.33 | 7.67 | 388.20 | 359.12 | 15.44 | 15.09 |
|  | 10.0 g L<sup>-1</sup> (K<sub>4</sub>) | 63.30 | 62.00 | 6.67 | 6.67 | 325.05 | 312.85 | 14.31 | 13.93 |
|  | Average | 68.86 | 66.27 | 7.13 | 7.67 | 385.83 | 376.63 | 15.15 | 14.96 |
| 40 mM (S<sub>1</sub>) | Control (K<sub>0</sub>) | 65.20 | 60.67 | 7.00 | 7.33 | 357.11 | 341.10 | 15.45 | 15.52 |
|  | 2.5 g L<sup>-1</sup> (K<sub>1</sub>) | 76.80 | 73.67 | 7.67 | 8.67 | 367.12 | 418.45 | 16.97 | 15.75 |
|  | 5.0 g L<sup>-1</sup> (K<sub>2</sub>) | 77.13 | 77.00 | 8.33 | 8.67 | 428.09 | 488.98 | 16.52 | 15.23 |
|  | 7.5 g L<sup>-1</sup> (K<sub>3</sub>) | 68.37 | 64.67 | 7.00 | 8.00 | 440.02 | 367.36 | 13.81 | 15.21 |
|  | 10.0 g L<sup>-1</sup> (K<sub>4</sub>) | 63.57 | 62.67 | 6.67 | 6.67 | 372.92 | 331.35 | 14.04 | 14.34 |
|  | Average | 70.21 | 67.73 | 7.33 | 7.87 | 393.05 | 389.45 | 15.36 | 15.21 |
| 80 mM (S<sub>2</sub>) | Control (K<sub>0</sub>) | 57.03 | 55.33 | 5.33 | 4.67 | 268.22 | 271.29 | 14.90 | 14.97 |
|  | 2.5 g L<sup>-1</sup> (K<sub>1</sub>) | 63.33 | 59.67 | 6.33 | 6.00 | 328.79 | 313.63 | 13.96 | 13.43 |
|  | 5.0 g L<sup>-1</sup> (K<sub>2</sub>) | 63.93 | 61.33 | 7.00 | 6.67 | 344.77 | 339.18 | 14.24 | 14.35 |
|  | 7.5 g L<sup>-1</sup> (K<sub>3</sub>) | 64.17 | 62.33 | 7.33 | 7.00 | 347.81 | 325.74 | 14.27 | 15.43 |
|  | 10.0 g L<sup>-1</sup> (K<sub>4</sub>) | 64.30 | 62.67 | 6.00 | 7.00 | 306.87 | 311.54 | 16.42 | 16.19 |
|  | Average | 62.55 | 60.27 | 6.40 | 6.27 | 319.29 | 312.27 | 14.76 | 14.88 |
| 120 mM (S<sub>3</sub>) | Control (K<sub>0</sub>) | 54.93 | 52.33 | 4.67 | 4.33 | 251.04 | 240.17 | 15.26 | 14.87 |
|  | 2.5 g L<sup>-1</sup> (K<sub>1</sub>) | 58.23 | 57.00 | 5.33 | 5.33 | 324.03 | 307.94 | 13.81 | 13.66 |
|  | 5.0 g L<sup>-1</sup> (K<sub>2</sub>) | 65.33 | 60.33 | 5.67 | 5.67 | 329.86 | 335.44 | 14.53 | 14.37 |
|  | 7.5 g L<sup>-1</sup> (K<sub>3</sub>) | 65.57 | 62.00 | 6.00 | 6.00 | 321.34 | 321.67 | 15.13 | 15.39 |
|  | 10.0 g L<sup>-1</sup> (K<sub>4</sub>) | 62.83 | 58.00 | 5.33 | 5.00 | 297.36 | 284.54 | 14.46 | 14.86 |
|  | Average | 61.38 | 57.93 | 5.40 | 5.27 | 304.72 | 297.95 | 14.64 | 14.63 |
| Averages across salinity level | Control (K<sub>0</sub>) | 60.79 | 57.92 | 5.92 | 5.92 | 307.93 | 300.80 | 15.26 | 15.14 |
|  | 2.5 g L<sup>-1</sup> (K<sub>1</sub>) | 68.13 | 64.92 | 6.67 | 7.08 | 354.13 | 358.38 | 15.01 | 14.51 |
|  | 5.0 g L<sup>-1</sup> (K<sub>2</sub>) | 70.18 | 67.58 | 7.17 | 7.33 | 390.92 | 407.66 | 15.14 | 14.83 |
|  | 7.5 g L<sup>-1</sup> (K<sub>3</sub>) | 66.17 | 63.50 | 6.92 | 7.17 | 374.34 | 343.47 | 14.66 | 15.28 |
|  | 10.0 g L<sup>-1</sup> (K<sub>4</sub>) | 63.50 | 61.33 | 6.17 | 6.33 | 325.55 | 310.07 | 14.81 | 14.83 |
| SEm ± (NaCl) |  | 1.29 | 1.12 | 0.27 | 0.33 | 4.11 | 3.99 | 0.35 | 0.45 |
| LSD (P=0.5), NaCl |  | 3.70 | 3.20 | 0.78 | 0.93 | 11.79 | 11.45 | NS | NS |
| SEm ± (KNO<sub>3</sub>) |  | 1.44 | 1.25 | 0.30 | 0.36 | 4.60 | 4.46 | 0.39 | 0.50 |
| LSD (P=0.5), KNO<sub>3</sub> |  | 4.14 | 3.58 | 0.87 | 1.04 | 13.19 | 12.81 | NS | NS |
| SEm ± (S× K) |  | 2.89 | 2.49 | 0.61 | 0.73 | 9.19 | 8.93 | 0.78 | 1.00 |
| LSD (P=0.5), S× K |  | NS | NS | NS | NS | 26.37 | 25.61 | NS | NS |

### 3.2. Leaf, stem and root biomass yield

Averaged across the KNO_3_ levels, yield (leaf, stem and root biomass) response to salinity levels varied significantly (*P* ≤ 0.05) as salinity increased up to NaCl at 120 mM (**Table 3**) during both growing seasons. Irrespective of KNO_3_ application, the maximum dry leaf yield of stevia (6.02 and 5.93 g plant^−1^) was found with the plant irrigated with low salinity water (NaCl at 40 mM) but remained on par (*P* ≥ 0.05) with control. In 1^st^ season, the reductions in dry leaf yield with high salinity level (NaCl at 120 mM) compared with control and low salinity (NaCl at 40 mM) were 23.80 and 26.07 %, respectively. Irrespective of foliar application of KNO_3_, plant irrigated with NaCl at 120 mM reduced root biomass significantly (*P* ≤ 0.05) compared with control by about 31 and 36 % in 1^st^ and 2^nd^ cropping season, respectively. Averaged across of salinity levels, yield (leaf, stem and root biomass) of stevia in response to foliar application of KNO_3_ increased significantly (*P* ≤ 0.05) as the concentration increased up to at 5.0 g L^−1^, thereafter decline trend was observed during both growing seasons (**Table 3**). The increases in dry leaf yield with KNO_3_ at 5.0 g L^−1^ over water spray as control were 26.59 and 33.04 % in 1^st^ and 2^nd^ years, respectively.

**Table 3.** Effect of salinity levels and foliar application of KNO_3_ on dry leaf yield, dry stem yield and dry root biomass of *Stevia rebaudiana*

| Salinity level (NaCl) | Concentration of KNO<sub>3</sub> | Dry leaf yield (g plant<sup>-1</sup>) |  | Dry stem yield (g plant<sup>-1</sup>) |  | Dry root biomass (g plant<sup>-1</sup>) |  |
| --- | --- | --- | --- | --- | --- | --- | --- |
|  |  | 2016 | 2017 | 2016 | 2017 | 2016 | 2017 |
| Control (S<sub>0</sub>) | Control (K<sub>0</sub>) | 5.47 | 5.33 | 6.35 | 5.82 | 1.84 | 2.05 |
|  | 2.5 g L<sup>-1</sup> (K<sub>1</sub>) | 6.07 | 5.99 | 6.67 | 6.90 | 1.87 | 2.10 |
|  | 5.0 g L<sup>-1</sup> (K<sub>2</sub>) | 7.02 | 7.18 | 7.36 | 8.21 | 2.15 | 2.38 |
|  | 7.5 g L<sup>-1</sup> (K<sub>3</sub>) | 5.99 | 5.39 | 6.96 | 6.58 | 2.06 | 2.09 |
|  | 10.0 g L<sup>-1</sup> (K<sub>4</sub>) | 4.66 | 4.37 | 5.67 | 5.21 | 1.64 | 1.51 |
|  | Average | 5.84 | 5.65 | 6.60 | 6.54 | 1.91 | 2.02 |
| 40 mM (S<sub>1</sub>) | Control (K<sub>0</sub>) | 5.50 | 5.30 | 6.23 | 5.72 | 1.72 | 1.77 |
|  | 2.5 g L<sup>-1</sup> (K<sub>1</sub>) | 6.21 | 6.55 | 7.29 | 7.65 | 2.16 | 2.41 |
|  | 5.0 g L<sup>-1</sup> (K<sub>2</sub>) | 7.07 | 7.45 | 7.89 | 8.64 | 2.46 | 2.61 |
|  | 7.5 g L<sup>-1</sup> (K<sub>3</sub>) | 6.09 | 5.57 | 6.95 | 6.76 | 1.84 | 2.03 |
|  | 10.0 g L<sup>-1</sup> (K<sub>4</sub>) | 5.24 | 4.75 | 6.73 | 5.58 | 1.51 | 1.36 |
|  | Average | 6.02 | 5.93 | 7.02 | 6.87 | 1.94 | 2.03 |
| 80 mM (S<sub>2</sub>) | Control (K<sub>0</sub>) | 4.01 | 4.06 | 4.99 | 4.83 | 1.20 | 1.14 |
|  | 2.5 g L<sup>-1</sup> (K<sub>1</sub>) | 4.59 | 4.22 | 5.84 | 5.36 | 1.39 | 1.22 |
|  | 5.0 g L<sup>-1</sup> (K<sub>2</sub>) | 4.92 | 4.86 | 6.38 | 6.15 | 1.66 | 1.61 |
|  | 7.5 g L<sup>-1</sup> (K<sub>3</sub>) | 4.96 | 5.01 | 6.44 | 6.35 | 1.96 | 2.08 |
|  | 10.0 g L<sup>-1</sup> (K<sub>4</sub>) | 5.02 | 5.05 | 5.50 | 5.77 | 1.99 | 2.09 |
|  | Average | 4.70 | 4.64 | 5.83 | 5.69 | 1.64 | 1.63 |
| 120 mM (S<sub>3</sub>) | Control (K<sub>0</sub>) | 3.83 | 3.57 | 4.75 | 4.22 | 1.09 | 1.09 |
|  | 2.5 g L<sup>-1</sup> (K<sub>1</sub>) | 4.48 | 4.21 | 5.11 | 4.76 | 1.32 | 1.25 |
|  | 5.0 g L<sup>-1</sup> (K<sub>2</sub>) | 4.79 | 4.81 | 5.63 | 5.65 | 1.49 | 1.29 |
|  | 7.5 g L<sup>-1</sup> (K<sub>3</sub>) | 4.86 | 4.95 | 5.70 | 5.73 | 1.62 | 1.69 |
|  | 10.0 g L<sup>-1</sup> (K<sub>4</sub>) | 4.31 | 4.21 | 4.75 | 4.56 | 1.08 | 1.11 |
|  | Average | 4.45 | 4.35 | 5.19 | 4.98 | 1.32 | 1.29 |
| Averages across salinity level | Control (K<sub>0</sub>) | 4.70 | 4.57 | 5.58 | 5.15 | 1.46 | 1.51 |
|  | 2.5 g L<sup>-1</sup> (K<sub>1</sub>) | 5.34 | 5.24 | 6.23 | 6.17 | 1.69 | 1.74 |
|  | 5.0 g L<sup>-1</sup> (K<sub>2</sub>) | 5.95 | 6.08 | 6.81 | 7.16 | 1.94 | 1.97 |
|  | 7.5 g L<sup>-1</sup> (K<sub>3</sub>) | 5.48 | 5.23 | 6.51 | 6.35 | 1.87 | 1.97 |
|  | 10.0 g L<sup>-1</sup> (K<sub>4</sub>) | 4.81 | 4.60 | 5.66 | 5.28 | 1.56 | 1.52 |
| SEm ± (NaCl) |  | 0.13 | 0.15 | 0.08 | 0.13 | 0.06 | 0.08 |
| LSD (P=0.5), NaCl |  | 0.37 | 0.42 | 0.24 | 0.38 | 0.18 | 0.24 |
| SEm ± (KNO<sub>3</sub>) |  | 0.14 | 0.17 | 0.09 | 0.15 | 0.07 | 0.09 |
| LSD (P=0.5), KNO<sub>3</sub> |  | 0.41 | 0.47 | 0.27 | 0.43 | 0.21 | 0.26 |
| SEm ± (S× K) |  | 0.29 | 0.33 | 0.19 | 0.30 | 0.15 | 0.18 |
| LSD (P=0.5), S× K |  | 0.82 | 0.95 | 0.53 | 0.86 | 0.41 | 0.53 |

Significant (*P* ≤ 0.05) interactions of salinity levels and KNO_3_ application on dry leaf and stem yield and root biomass occurred in both seasons (**Table 3**). At control (irrigated with normal water) and low salinity level (NaCl at 40 mM), dry leaf, stem and root biomass increased as concentration of KNO_3_ increased from 0 to 5.0 g L^−1^, while the reverse trend was observed with further increases in KNO_3_ concentration. However, at moderate salinity level (NaCl at 80 mM), dry leaf yield was increased with corresponding increase in concentration of KNO_3_ up to 10.0 g L^−1^ (**Table 3**).

### 3.3. Chlorophyll (Chl) concentration in leaf

The photosynthetic pigments (Chl_*a*_, Chl_*b*_, total Chl_*a+b*_, and Chl_*a*_: Chl_*b*_) in stevia leaf as influenced by saline levels and foliar application of KNO_3_ are presented in **Fig 1a-d**. Averaged over KNO_3_ level, there were significant (*P* ≤ 0.05) differences among the salinity levels in concentration of Chl_*a*_, Chl_*b*_, total, and Chl, during both years. The concentrations of photosynthetic pigments were gradually decreased at higher salinity level, irrespective of foliar spray. Averaged across the salinity levels, the effects of foliar application of KNO_3_ on Chl_*a*_, Chl_*b*_ and total Chl content in leaves were significant (*P* ≤ 0.05) in both years. The Chl_*a*_ and total Chl contents in leaves were increased with the increasing concentration of KNO_3_ up to 5.0 g L^−1^ in both seasons (**Fig. 1a, c**)..

**Fig.1.**
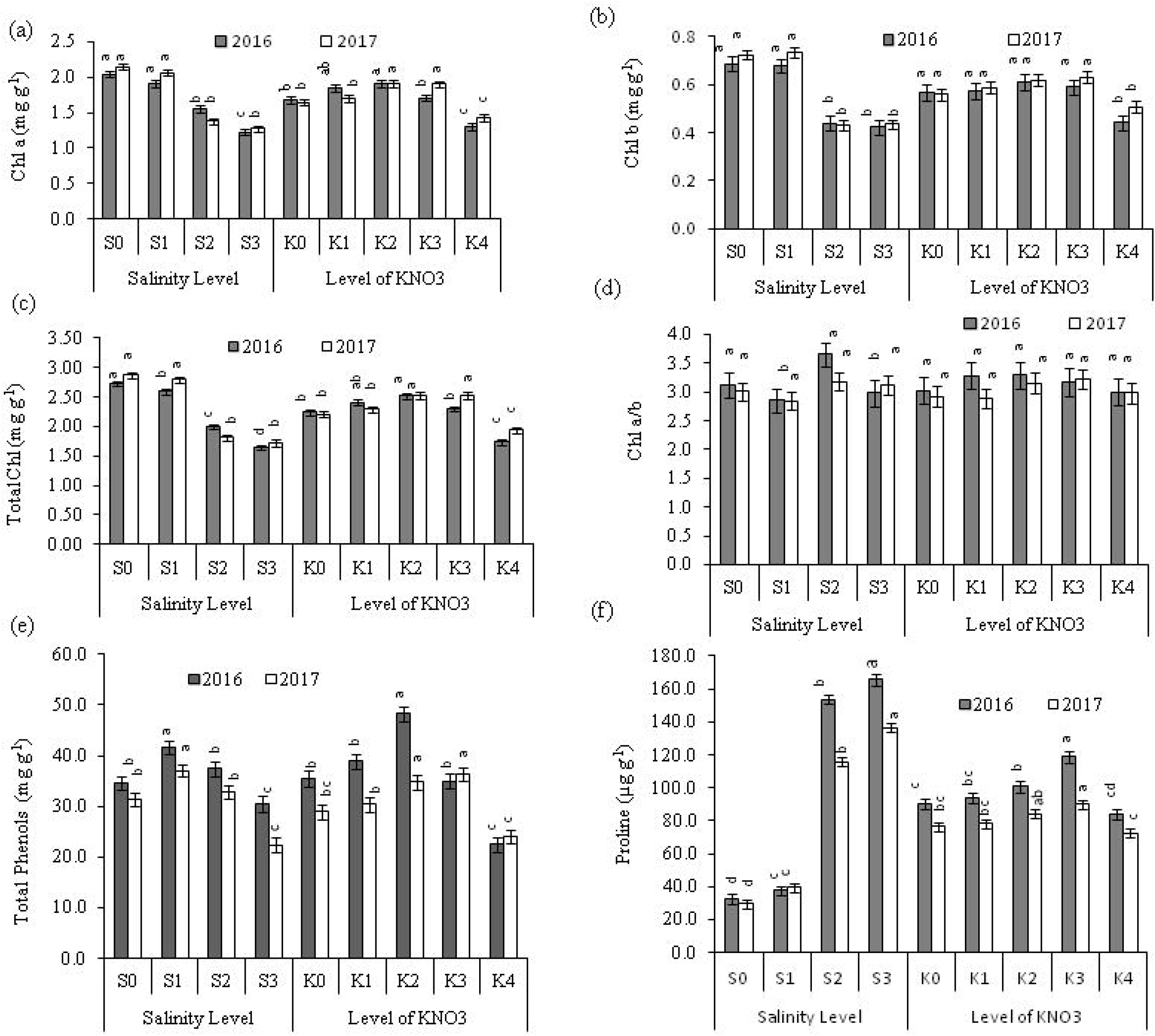
Variation in biochemical parameters i.e. (a) chlorophyll a, (b) chlorophyll b, (c) total chlorophyll, (d) chl a/ chl b, (e) total phenols and (f) proline in stevia leaves under different salinity and KNO_3_ levels. S_0_, S_1_, S_2_, and S_3_ are representing salinity level (NaCl) of 0.0, 40, 80 and 120 mM, respectively, whereas as K_0_, K_1_, K_2_, K_3_, K_4_ are representing the folia application of KNO_3_ at 0.0, 2.5, 5.0, 7.5, and 10.0 g L^−1^, respectively. Vertical bars represented the mean standard errors (±).

The interaction effects between salinity levels and foliar application of KNO_3_ on Chl_*a*_ and total Chl content in leaves were found significant (*P* ≤ 0.05) in both years (**Table 4**). At low salinity levels (control and NaCl at 40 mM), although total Chl contents were higher with foliar application of KNO_3_ at 5.0 g L^−1^ than in water spray control in 2^nd^ year, but these differences were statistically insignificant (*P* ≥ 0.05). On the other hand, the foliar application of KNO_3_ at 5.0 g L^−1^ registered significantly (*P* ≤ 0.05) higher total Chl content in leaves compared with water spray control (**Table 4**) at higher salinity level (NaCl at 80 and 120 mM).

**Table 4.** Interaction effects between salinity level and application of KNO_3_ on biochemical parameters and nutrient accumulation in leaf of *Stevia rebaudiana*

| Salinity levels (NaCl) | Concentration of KNO<sub>3</sub> | Biochemical parameters |  |  |  |  |  | Ion accumulation in leaf |  |  |  |
| --- | --- | --- | --- | --- | --- | --- | --- | --- | --- | --- | --- |
|  |  | Total Chl (mg g<sup>-1</sup>) |  | Proline (μg g<sup>-1</sup>) |  | Total Phenols (mg g<sup>-1</sup>) |  | Na<sup>+</sup> (mg g<sup>-1</sup>) |  | K<sup>+</sup> (mg g<sup>-1</sup>) |  |
|  |  | 2016 | 2017 | 2016 | 2017 | 2016 | 2017 | 2016 | 2017 | 2016 | 2017 |
| Control (S<sub>0</sub>) | Control (K<sub>0</sub>) | 2.73 | 2.94 | 23.48 | 29.07 | 32.25 | 28.79 | 1.30 | 1.00 | 22.32 | 22.80 |
|  | 2.5 g L<sup>-1</sup> (K<sub>1</sub>) | 2.90 | 2.97 | 26.57 | 29.30 | 39.58 | 33.92 | 1.13 | 1.05 | 23.05 | 24.20 |
|  | 5.0 g L<sup>-1</sup> (K<sub>2</sub>) | 2.93 | 3.12 | 27.90 | 30.89 | 46.21 | 35.00 | 1.07 | 1.00 | 25.23 | 25.57 |
|  | 7.5 g L<sup>-1</sup> (K<sub>3</sub>) | 2.64 | 2.83 | 36.48 | 29.15 | 29.88 | 29.83 | 0.90 | 0.92 | 23.53 | 24.40 |
|  | 10.0 g L<sup>-1</sup> (K<sub>4</sub>) | 2.48 | 2.51 | 49.29 | 28.99 | 25.50 | 29.38 | 0.68 | 0.92 | 21.33 | 23.32 |
| 40 mM (S<sub>1</sub>) | Control (K<sub>0</sub>) | 2.81 | 2.69 | 36.10 | 38.26 | 38.04 | 28.13 | 1.57 | 1.32 | 23.92 | 23.28 |
|  | 2.5 g L<sup>-1</sup> (K<sub>1</sub>) | 2.89 | 2.83 | 34.14 | 36.98 | 42.42 | 31.38 | 1.15 | 1.07 | 24.62 | 25.02 |
|  | 5.0 g L<sup>-1</sup> (K<sub>2</sub>) | 2.79 | 2.87 | 35.10 | 39.26 | 68.17 | 43.38 | 1.15 | 1.02 | 26.63 | 26.98 |
|  | 7.5 g L<sup>-1</sup> (K<sub>3</sub>) | 2.63 | 2.91 | 37.52 | 41.32 | 41.21 | 54.04 | 1.03 | 0.93 | 24.85 | 25.62 |
|  | 10.0 g L<sup>-1</sup> (K<sub>4</sub>) | 1.83 | 2.72 | 45.14 | 41.63 | 17.83 | 28.42 | 0.85 | 0.95 | 21.97 | 22.83 |
| 80 mM (S<sub>2</sub>) | Control (K<sub>0</sub>) | 1.94 | 1.66 | 122.90 | 111.38 | 39.50 | 37.63 | 1.62 | 1.48 | 16.22 | 16.20 |
|  | 2.5 g L<sup>-1</sup> (K<sub>1</sub>) | 2.12 | 1.70 | 164.10 | 113.47 | 40.21 | 35.38 | 1.52 | 1.28 | 17.98 | 18.45 |
|  | 5.0 g L<sup>-1</sup> (K<sub>2</sub>) | 2.43 | 2.06 | 165.10 | 111.36 | 45.79 | 36.79 | 1.23 | 1.17 | 18.67 | 19.08 |
|  | 7.5 g L<sup>-1</sup> (K<sub>3</sub>) | 2.15 | 2.28 | 198.33 | 136.78 | 37.08 | 35.96 | 1.50 | 1.10 | 18.67 | 19.77 |
|  | 10.0 g L<sup>-1</sup> (K<sub>4</sub>) | 1.33 | 1.36 | 118.43 | 106.47 | 24.38 | 18.54 | 0.97 | 1.05 | 18.05 | 19.65 |
| 120 mM (S<sub>3</sub>) | Control (K<sub>0</sub>) | 1.49 | 1.55 | 177.76 | 127.79 | 32.00 | 21.33 | 2.07 | 1.62 | 15.50 | 12.53 |
|  | 2.5 g L<sup>-1</sup> (K<sub>1</sub>) | 1.74 | 1.66 | 150.00 | 133.45 | 33.29 | 20.75 | 1.55 | 1.37 | 14.43 | 15.18 |
|  | 5.0 g L<sup>-1</sup> (K<sub>2</sub>) | 1.92 | 2.08 | 174.81 | 155.27 | 33.00 | 24.17 | 1.35 | 1.30 | 16.77 | 16.50 |
|  | 7.5 g L<sup>-1</sup> (K<sub>3</sub>) | 1.79 | 2.12 | 203.16 | 146.32 | 31.92 | 25.58 | 1.22 | 1.23 | 15.07 | 15.98 |
|  | 10.0 g L<sup>-1</sup> (K<sub>4</sub>) | 1.32 | 1.16 | 122.33 | 112.56 | 22.29 | 19.83 | 0.95 | 1.20 | 13.17 | 13.48 |
| SEm± for (S× K) |  | 0.09 | 0.11 | 6.75 | 5.46 | 3.02 | 2.88 | 0.07 | 0.04 | 0.33 | 0.48 |
| CD (P=0.5) |  | 0.26 | 0.32 | 19.36 | 15.67 | 8.66 | 8.25 | 0.19 | 0.12 | 0.95 | 1.39 |

### 3.4. Accumulation of total phenols and proline in leaf

Averaged over the KNO_3_ levels, at 40 mM NaCl, total phenols (41.53 and 37.07 mg g^−1^) content in leaves was 19.75 and 15.35 % higher than that of control (34.68 and 31.38 mg g^−1^) in 1^st^ and 2^nd^ cropping seasons, respectively (**Fig. 1e**). It has also been observed that a significant (*P* ≤ 0.05) decrease in total phenols content in leaves was occurred in parallel with further increase in NaCl concentration, and about 27 and 40 % decreases were registered in the plants subjected to NaCl at 120 mM compared with the plants treated with NaCl at 40 mM in 1^st^ and 2^nd^ cropping seasons, respectively. Averaged over salinity level, the maximum value (48.29 mg g^−1^) was observed with KNO_3_ at 5.0 g L^−1^, which was 36.22 % higher than control in 2016. In 2017, by increasing the concentration of KNO_3_ from control to 7.5 g L^−1^, the concentration of total phenols was increased from 28.97 to 36.35 mg g^−1^. As NaCl concentration was increased, in parallel accumulation of proline content was increased (**Fig. 2f**). Averaged across salinity level, there was a significant (*P* ≤ 0.05) difference in proline content between the leaves treated with water spray and higher concentration of KNO_3_ (at 5.0 and 7.5 g L^−1^). At 7.5 g L^−1^ KNO_3_, 31.99 % and 17.54 % increases were observed in proline content compared with control in 2016 and 2017, respectively. However, proline content was sharply decreased with KNO_3_ at 10.0 g L^−1^ (**Fig. 1f**).

**Fig. 2.**
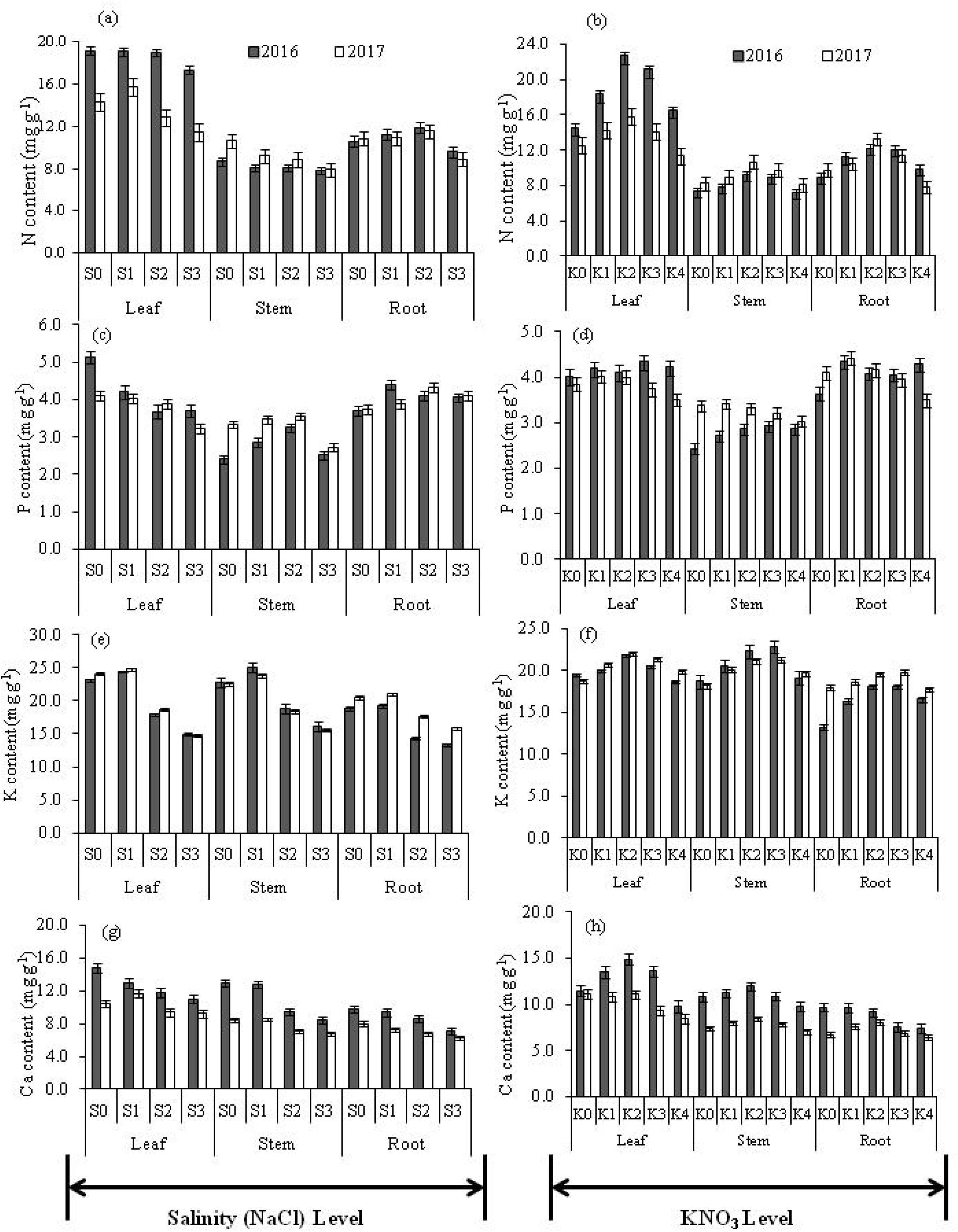
Effects of salinity level and exogenous application of KNO_3_ on accumulation of total nitrogen (a, b) phosphorus (c, d), potassium (e, f), and calcium (g, h), in the leaf, stem and root of *Stevia rebaudiana.* S_0_, S_1_, S_2_, and S_3_ represents as control, NaCl at 40, 80, and 120 mM, respectively. K_0_, K_1_, K_2_, K_3_, K_4_ are representing the folia application of KNO_3_ at 0.0, 2.5, 5.0, 7.5, and 10.0 g L^−1^, respectively. The mean standard errors (±) are presented with vertical bars.

Significant (*P* ≤ 0.05) interaction effects between salinity level and KNO_3_ on total phenols and proline content in stevia leaves were occurred in both years (**Table 4**). In 1^st^ season, total phenols content was significantly increased with KNO_3_ at 5.0 g L^−1^ compared with water spray and higher concentration of KNO_3_ (10.0 g L^−1^) under non-saline (control) and low saline (NaCl at 40 mM) conditions. Proline content was not significantly (*P* ≥ 0.05) influenced by the foliar application of KNO_3_ at low salinity level (NaCl at 40 mM), whereas significantly (*P* ≤ 0.05) higher amount was recorded with KNO_3_ at 10.0 g L^−1^ compared with water spray under non-saline condition in 1^st^ year. On the other hand, at moderate salinity level (NaCl at 80 mM), proline content was increased with the corresponding increase in concentration of KNO_3_ up to 7.5 g L^−1^ (**Table 4**).

### 3.5. Ions distribution in leaf, stem and root

At harvest stage, ions (N, P, K^+^, Ca^2+^, Na^+^ and Cl^−^) accumulation patterns in leaf, stem and root as influenced by different salinity levels and foliar applications of KNO_3_ are presented in **Fig. 2 and 3**. Averaged over KNO_3_ level, concentration of N in leaf was significantly (*P* ≤ 0.05) decreased from 19.18 and 14.37 mg g^−1^ to 17.36 and 11.8 mg g^−1^ at higher salinity level (NaCl at 120 mM) in 2016 and 2017 cropping seasons, respectively (**Fig. 2a**). In root, significantly (*P* ≤ 0.05) higher amount (11.90 and 11.53 mg g^−1^) of N was recorded with plants treated with NaCl at 80 mM compared with NaCl at 120 mM. Regardless of KNO_3_ level, the maximum K^+^ contents in leaf (24.40 and 24.75 mg g^−1^), stem (25.04 and 23.74 mg g^−1^), and root (19.22 and 21.02 mg g^−1^) were registered with the plants irrigated with low salinity water (NaCl at 40 mM); however, sharply decreased with NaCl at 120 mM during both growing seasons(**Fig. 2e**). The Ca^2+^ uptake was also reduced significantly (*P* ≤ 0.05) in the plants treated with high concentration of NaCl (120 mM) (**Fig. 2g**).

Averaged over KNO_3_ level, the effects of salinity level on the accumulation of Na^+^ and Cl^−^ in leaf, stem and root were found to be significant (*P* ≤ 0.05) in both years (**Fig. 3a,c**). As NaCl concentration was increased, in parallel accumulations of Na^+^ and Cl^−^ in leaf, stem and root were increased. Averaged across KNO_3_ level the maximum and lowest values of K^+^: Na^+^ and Ca^2+^: Na^+^ in all parts (except K^+^: Na^+^ in stem in 2016 and Ca^2+^: Na^+^ in leaf in 2017) were registered with control (irrigated with non-saline water) and NaCl at 120 mM, respectively (**Fig. 3e, g**).

**Fig. 3.**
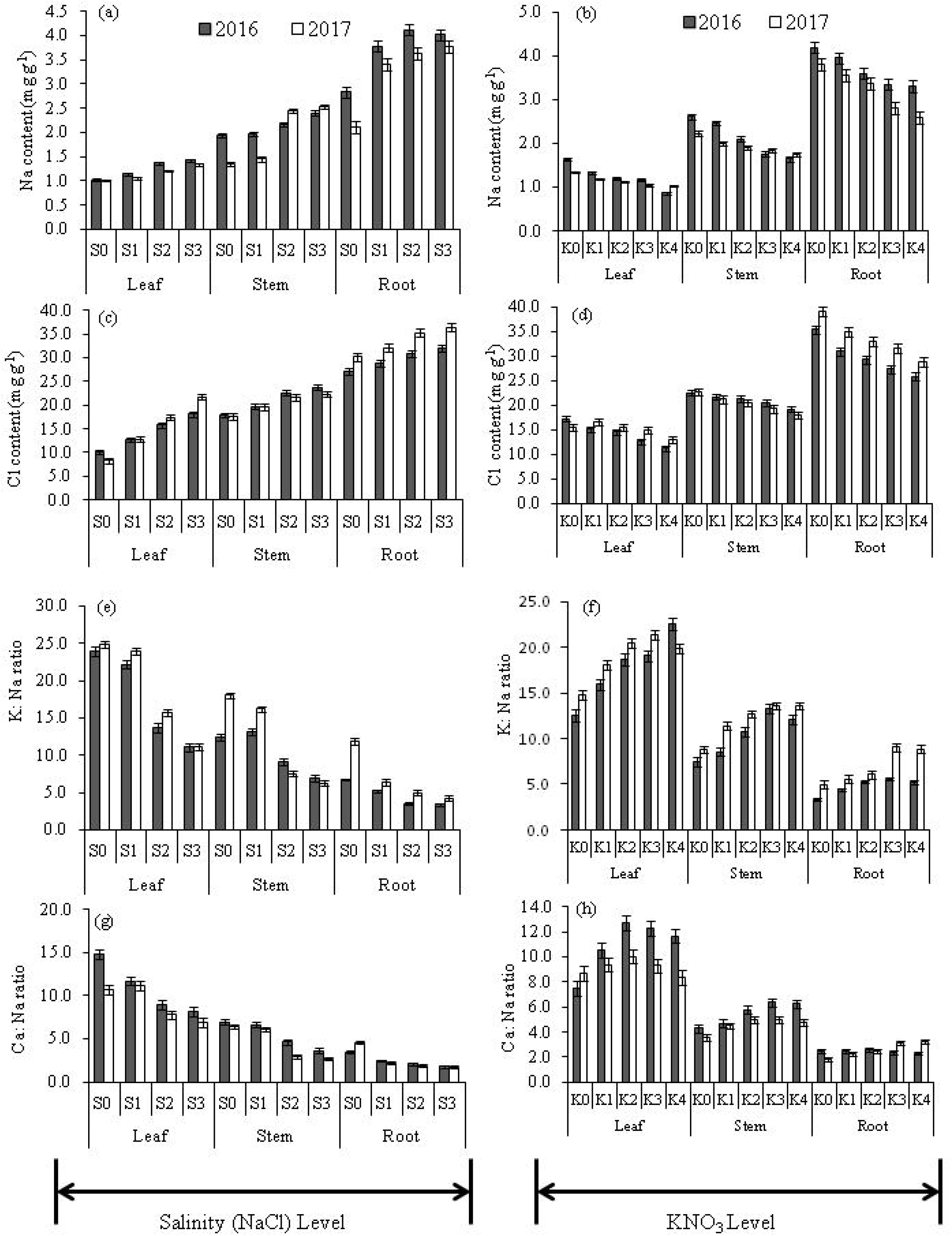
Effects of salinity level and exogenous application of KNO_3_ on accumulation of sodium (a, b) chlorine (c, d), K: Na (e, f), and Ca: Na (g, h), in the leaf, stem and root of *Stevia rebaudiana.* S_0_, S_1_, S_2_, and S_3_ represents as control, NaCl at 40, 80, and 120 mM, respectively. K_0_, K_1_, K_2_, K_3_, K_4_ are representing the foliar application of KNO_3_ at 0.0, 2.5, 5.0, 7.5, and 10.0 g L^−1^, respectively. The mean standard errors (±) are presented with vertical bars.

On the other hand, pooled across salinity level, the accumulation patterns of N and Ca^2+^ in leaf, stem and root were gradually increased with increasing the level of KNO_3_ concentration up to 5.0 g L^−1^, thereafter decline trend was observed in both the cropping seasons (**Fig. 2b, h**). On the other hand, pooled across all salinity environments, accumulation of K in leaf was significantly (*P* ≤ 0.05) higher with KNO_3_ at 5.0 g L^−1^ compared with rest of the treatments in both seasons (**Fig. 2f**). Pooled across the salinity level, accumulations of Na^+^ and Cl^−^ in all plant organs were sharply declined with corresponding increases of KNO_3_ level (**Fig. 3b, d**). On the other hand, the ratios of K^+^: Na^+^ in leaf, stem and root were influenced significantly (*P* ≤ 0.05) by the foliar application of KNO_3_, and the lowest values were registered with water spray as control, which were significantly (*P* ≤ 0.05) different from the application of KNO_3_ at 5.0 g L^−1^ and higher concentration (**Fig. 3f**). In some cases (particularly K^+^ and Na^+^ in leaf), the significant (*P* ≤ 0.05) interaction effects between salinity and KNO_3_ level on ions accumulation were also observed (**Table 4**).

### 3.6. Accumulation of Steviol glycosides (SGs)

Total eight SGs namely stevioside, Reb-A, Reb-F, Reb-C, dulcoside-A, rubusoside, Reb-B, and steviolbioside were analyzed in this experiment to understand the influence of salinity stress and foliar application of KNO_3_. Here, most two prominent SGs, Stevioside and Reb-A, and total SGs are presented in **Fig. 4a, b**. In case of total steviol glycosides (TSGs), the sum of eight aforesaid mentioned SGs has been considered. The plants treated with low salinity water (NaCl at 40 mM) accumulated significantly (*P*≤0.05) higher amount of stevioside (48.43 and 39.35 mg g^−1^ dry leaf) compared with the plants treated with moderate (NaCl at 80 mM) and high salinity (NaCl at 120 mM) water in both years (**Fig. 4a**). Reb-A content was also significantly (*P*<0.05) influenced by salinity level, and high salinity level (NaCl at 120 mM) produced significant lower Reb-A (17.38 and 18.34 mg g^−1^) compared with non-salinity treatment in both years (**Fig.4b**). In case of TSGs yield (g plant^−1^), there was no significant (*P* ≥ 0.05) difference between low saline (NaCl at 40 mM) water treated plants and non-saline water treated plants in 1^st^ and 2^nd^ cropping season (**Fig.4d**).

**Fig. 4.**
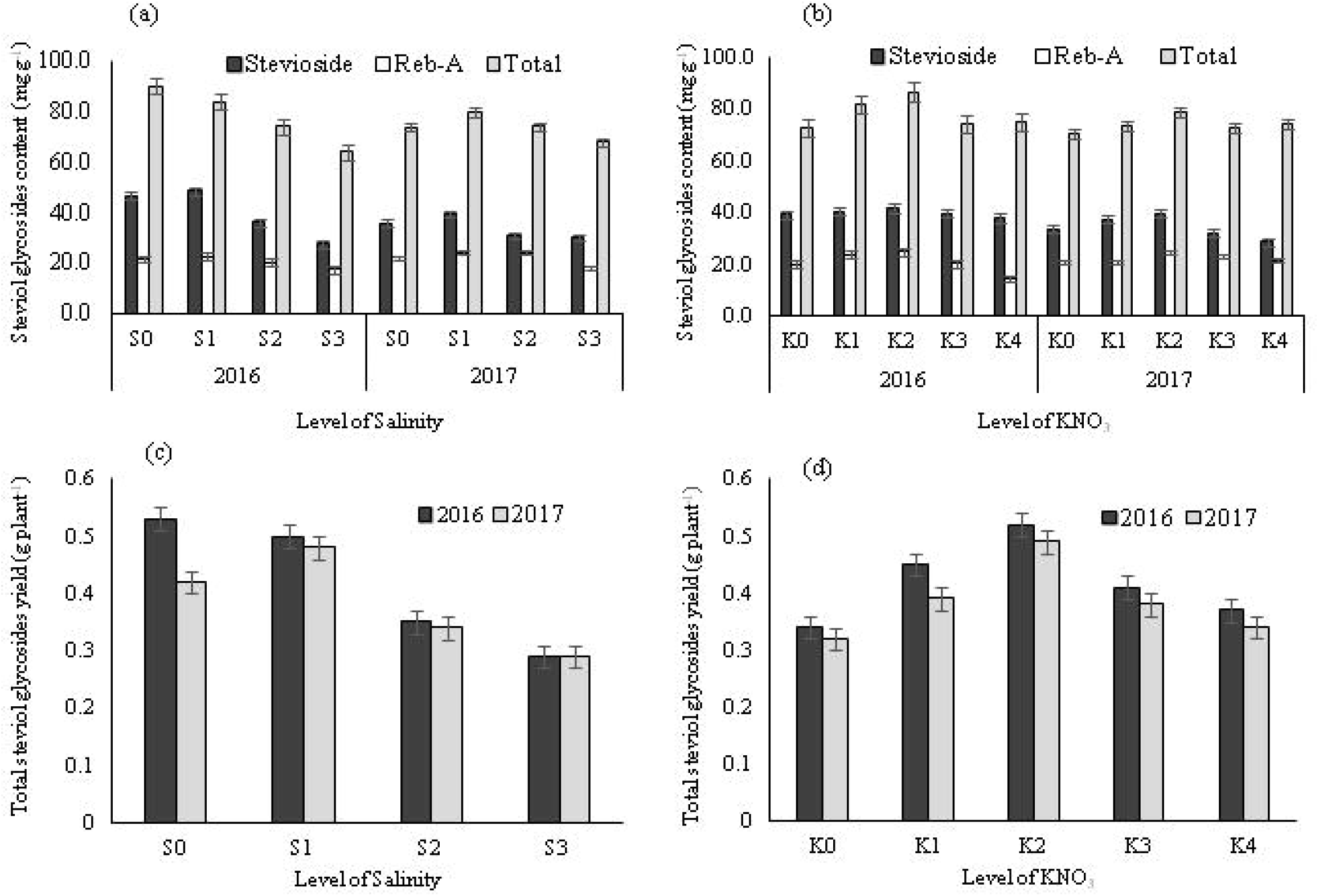
Steviol glycosides accumulation (mg g^−1^) in leaf as influenced by salinity level and folia application of KNO_3_ (a, b). Steviol glycosides yield are represented in g plant^−1^ (c,d). S_0_, S_1_, S_2_, and S_3_ are representing salinity level (NaCl) of 0.0, 40, 80 and 120 mM, respectively, whereas as K_0_, K_1_, K_2_, K_3_, K_4_ are representing the folia application of KNO_3_ at 0.0, 2.5, 5.0, 7.5, and 10.0 g L^−1^, respectively. The mean standard errors (±) are presented with vertical bars.

Averaged over salinity levels, there were no significant (*P*≤0.05) differences among the KNO_3_ levels for accumulation of stevioside in 1^st^ year. However, significant (*P* ≤ 0.05) differences among the KNO_3_ levels were observed in 2^nd^ year, and the maximum (39.58 mg g^−1^) value was recorded with KNO_3_ at 5.0 g L^−1^ (**Fig.4b**). Irrespective of salinity levels, the maximum quantity of Reb-A (24.75 and 24.80 mg g^−1^) was also recorded with KNO_3_ at 5.0 g L^−1^, which was significantly (P ≤ 0.05) higher than control during both years. Significant (*P*≤0.05) variations in the TSGs yield (g plant^−1^) in response to different KNO_3_ levels were found during both years, and the maximum yield (0.52 and 0.49 g plant^−1^) was recorded with the application of KNO_3_ at 5.0 g L^−1^ (**Fig. 4d**).

The interaction effects between salinity levels and foliar applications of KNO_3_ on accumulation of stevioside, Reb-A and TSGs and TSGs yield (g plant^−1^) were found to be significant (*P* ≤ 0.05) in both seasons (**Table 5**). Effect of KNO_3_ on accumulation of stevioside was significant (*P* ≤ 0.05) under non-saline condition (control) in both years, and the maximum value was recorded with KNO_3_ at 5.0 g L^−1^ (62.85 and 43.72 mg g^−1^). Under other salinity levels, accumulation patterns of stevioside due to foliar application of KNO_3_ were found to be irregular.

**Table 5.**
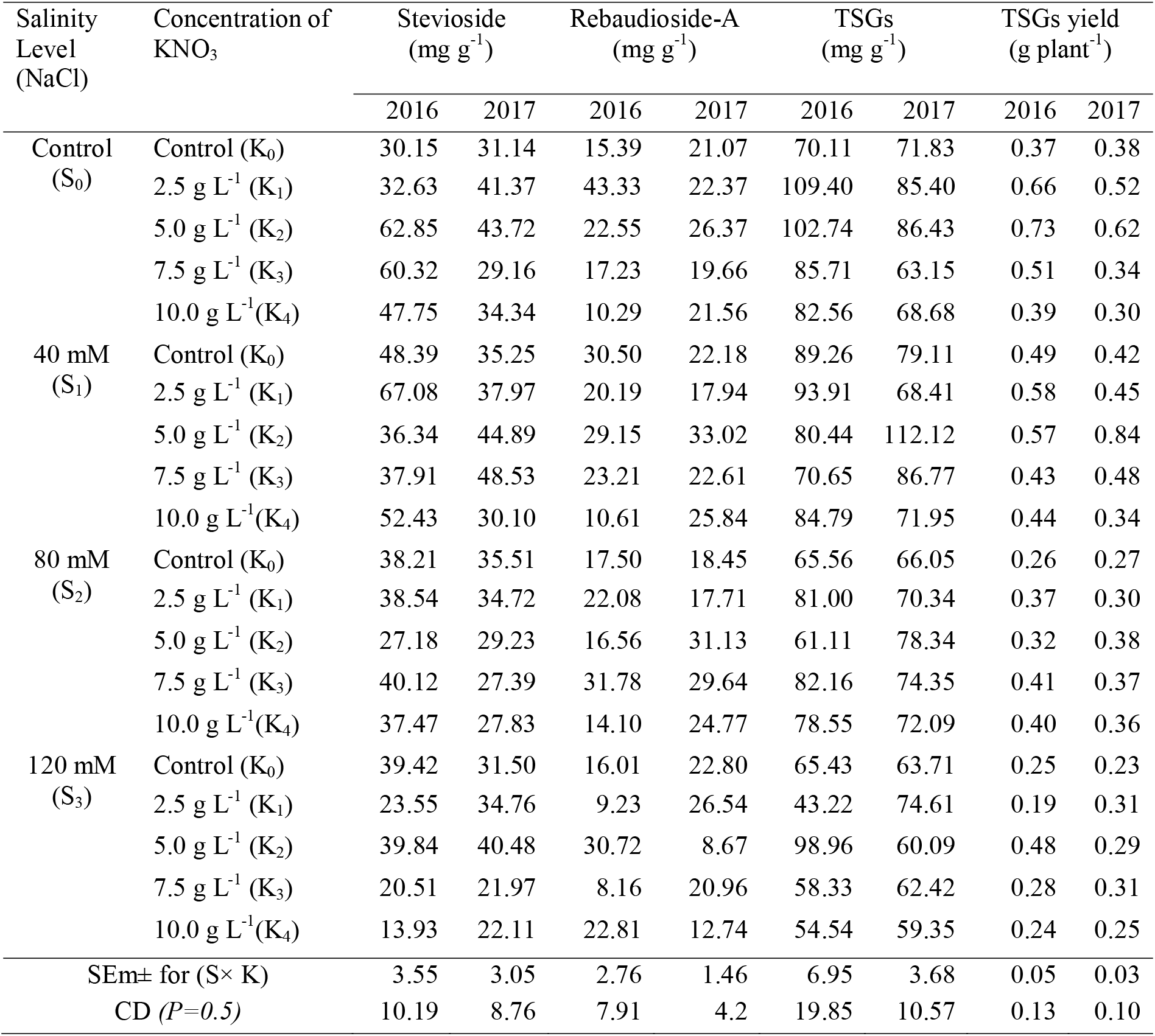
The interaction effects between salinity levels and foliar application of KNO_3_ on secondary metabolites profile in *Stevia rebaudiana*

| Salinity Level (NaCl) | Concentration of KNO<sub>3</sub> | Stevioside (mg g<sup>-1</sup>) |  | Rebaudioside-A (mg g<sup>-1</sup>) |  | TSGs (mg g<sup>-1</sup>) |  | TSGs yield (g plant<sup>-1</sup>) |  |
| --- | --- | --- | --- | --- | --- | --- | --- | --- | --- |
|  |  | 2016 | 2017 | 2016 | 2017 | 2016 | 2017 | 2016 | 2017 |
| Control (S<sub>0</sub>) | Control (K<sub>0</sub>) | 30.15 | 31.14 | 15.39 | 21.07 | 70.11 | 71.83 | 0.37 | 0.38 |
|  | 2.5 g L<sup>-1</sup> (K<sub>1</sub>) | 32.63 | 41.37 | 43.33 | 22.37 | 109.40 | 85.40 | 0.66 | 0.52 |
|  | 5.0 g L<sup>-1</sup> (K<sub>2</sub>) | 62.85 | 43.72 | 22.55 | 26.37 | 102.74 | 86.43 | 0.73 | 0.62 |
|  | 7.5 g L<sup>-1</sup> (K<sub>3</sub>) | 60.32 | 29.16 | 17.23 | 19.66 | 85.71 | 63.15 | 0.51 | 0.34 |
|  | 10.0 g L<sup>-1</sup> (K<sub>4</sub>) | 47.75 | 34.34 | 10.29 | 21.56 | 82.56 | 68.68 | 0.39 | 0.30 |
| 40 mM (S<sub>1</sub>) | Control (K<sub>0</sub>) | 48.39 | 35.25 | 30.50 | 22.18 | 89.26 | 79.11 | 0.49 | 0.42 |
|  | 2.5 g L<sup>-1</sup> (K<sub>1</sub>) | 67.08 | 37.97 | 20.19 | 17.94 | 93.91 | 68.41 | 0.58 | 0.45 |
|  | 5.0 g L<sup>-1</sup> (K<sub>2</sub>) | 36.34 | 44.89 | 29.15 | 33.02 | 80.44 | 112.12 | 0.57 | 0.84 |
|  | 7.5 g L<sup>-1</sup> (K<sub>3</sub>) | 37.91 | 48.53 | 23.21 | 22.61 | 70.65 | 86.77 | 0.43 | 0.48 |
|  | 10.0 g L<sup>-1</sup> (K<sub>4</sub>) | 52.43 | 30.10 | 10.61 | 25.84 | 84.79 | 71.95 | 0.44 | 0.34 |
| 80 mM (S<sub>2</sub>) | Control (K<sub>0</sub>) | 38.21 | 35.51 | 17.50 | 18.45 | 65.56 | 66.05 | 0.26 | 0.27 |
|  | 2.5 g L<sup>-1</sup> (K<sub>1</sub>) | 38.54 | 34.72 | 22.08 | 17.71 | 81.00 | 70.34 | 0.37 | 0.30 |
|  | 5.0 g L<sup>-1</sup> (K<sub>2</sub>) | 27.18 | 29.23 | 16.56 | 31.13 | 61.11 | 78.34 | 0.32 | 0.38 |
|  | 7.5 g L<sup>-1</sup> (K<sub>3</sub>) | 40.12 | 27.39 | 31.78 | 29.64 | 82.16 | 74.35 | 0.41 | 0.37 |
|  | 10.0 g L<sup>-1</sup> (K<sub>4</sub>) | 37.47 | 27.83 | 14.10 | 24.77 | 78.55 | 72.09 | 0.40 | 0.36 |
| 120 mM (S<sub>3</sub>) | Control (K<sub>0</sub>) | 39.42 | 31.50 | 16.01 | 22.80 | 65.43 | 63.71 | 0.25 | 0.23 |
|  | 2.5 g L<sup>-1</sup> (K<sub>1</sub>) | 23.55 | 34.76 | 9.23 | 26.54 | 43.22 | 74.61 | 0.19 | 0.31 |
|  | 5.0 g L<sup>-1</sup> (K<sub>2</sub>) | 39.84 | 40.48 | 30.72 | 8.67 | 98.96 | 60.09 | 0.48 | 0.29 |
|  | 7.5 g L<sup>-1</sup> (K<sub>3</sub>) | 20.51 | 21.97 | 8.16 | 20.96 | 58.33 | 62.42 | 0.28 | 0.31 |
|  | 10.0 g L<sup>-1</sup> (K<sub>4</sub>) | 13.93 | 22.11 | 22.81 | 12.74 | 54.54 | 59.35 | 0.24 | 0.25 |
| SEm± for (S× K) |  | 3.55 | 3.05 | 2.76 | 1.46 | 6.95 | 3.68 | 0.05 | 0.03 |
| CD (P=0.5) |  | 10.19 | 8.76 | 7.91 | 4.2 | 19.85 | 10.57 | 0.13 | 0.10 |

### 3.7. Correlation, regression and principle component analysis

The statistical relationships among the agronomic traits (Plant height, number of branch, LA, SLW, dry root, stem and leaf yield) were presented in a correlation matrix (**Table S1**). The data revealed that most of the agronomic traits were positively correlated with each other. The relationship of dry leaf yield of stevia with the concentration of KNO_3_ was best described by the second-degree curve (**Fig.5**) with regression equations as given below:

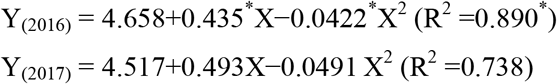

**Fig. 5.**
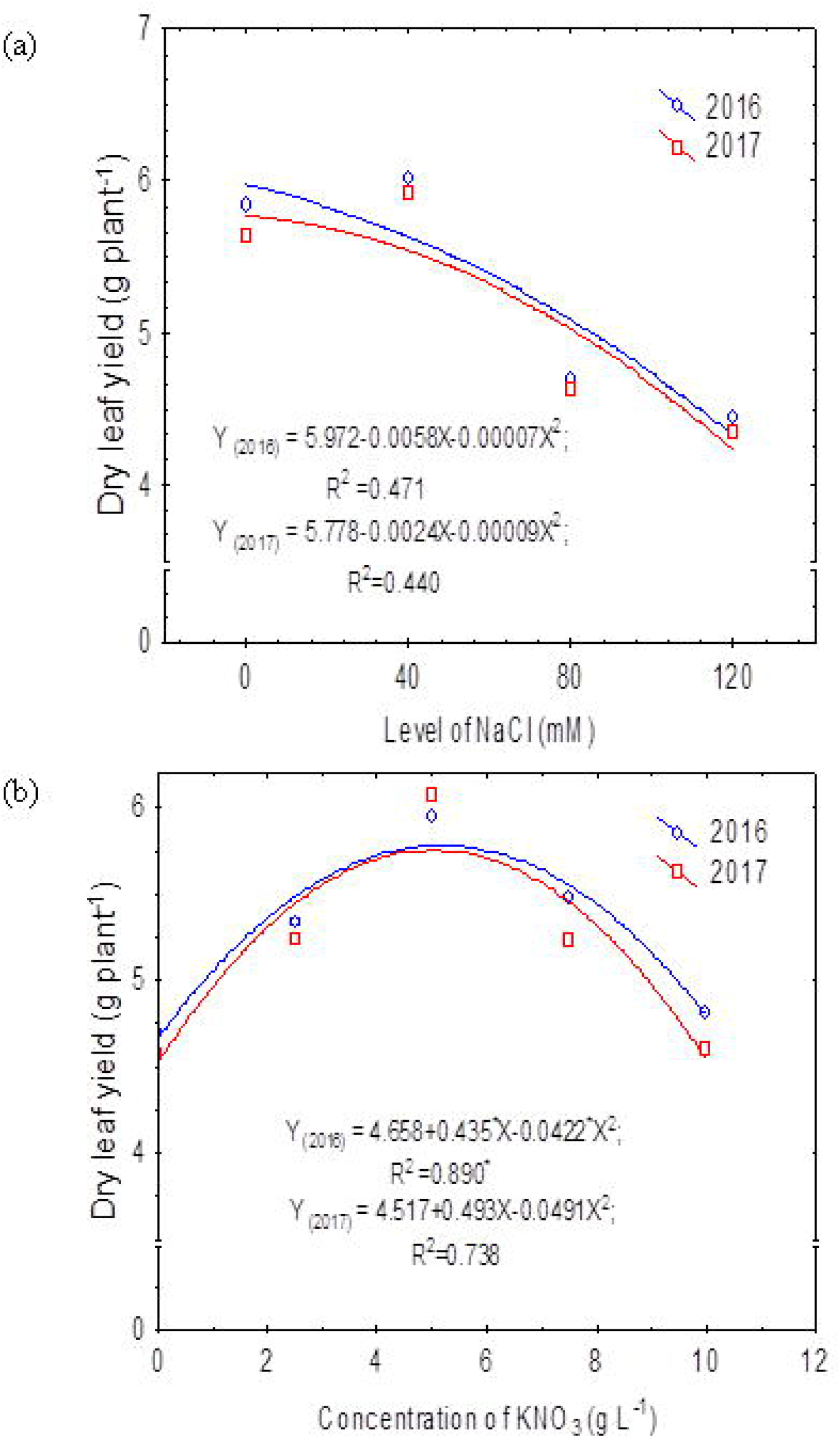
Regression equation between dry leaf yield and salinity level (a). Another regression equation between dry leaf yield and concentration of KNO_3_ (b).

The dry leaf yield was increased with the corresponding increase in concentration of KNO_3_ up to 5.0 g L^−1^, and thereafter trend was declined in both the years.

In this investigation, the principle component analysis (PCA) was conducted on six agronomic traits (plant height, branch, leaf area, dry root yield, dry leaf yield, dry stem yield) and eleven chemical and biochemical parameters (total Chl, total phenols, proline, N, P, K^+^, Ca^2+^, Na^+^, Cl^−^, K: Na, Ca: Na) of stevia to understand the interaction effects of treatment combination on these traits and their relationship (**Fig. 6 a-d**). The data revealed that the first two principal components (PC_1_ and PC_2_) were cumulatively accounted for 80.32% and 80.07 % of the total variations for the 1^st^ and 2^nd^ year, respectively.

**Fig. 6.**
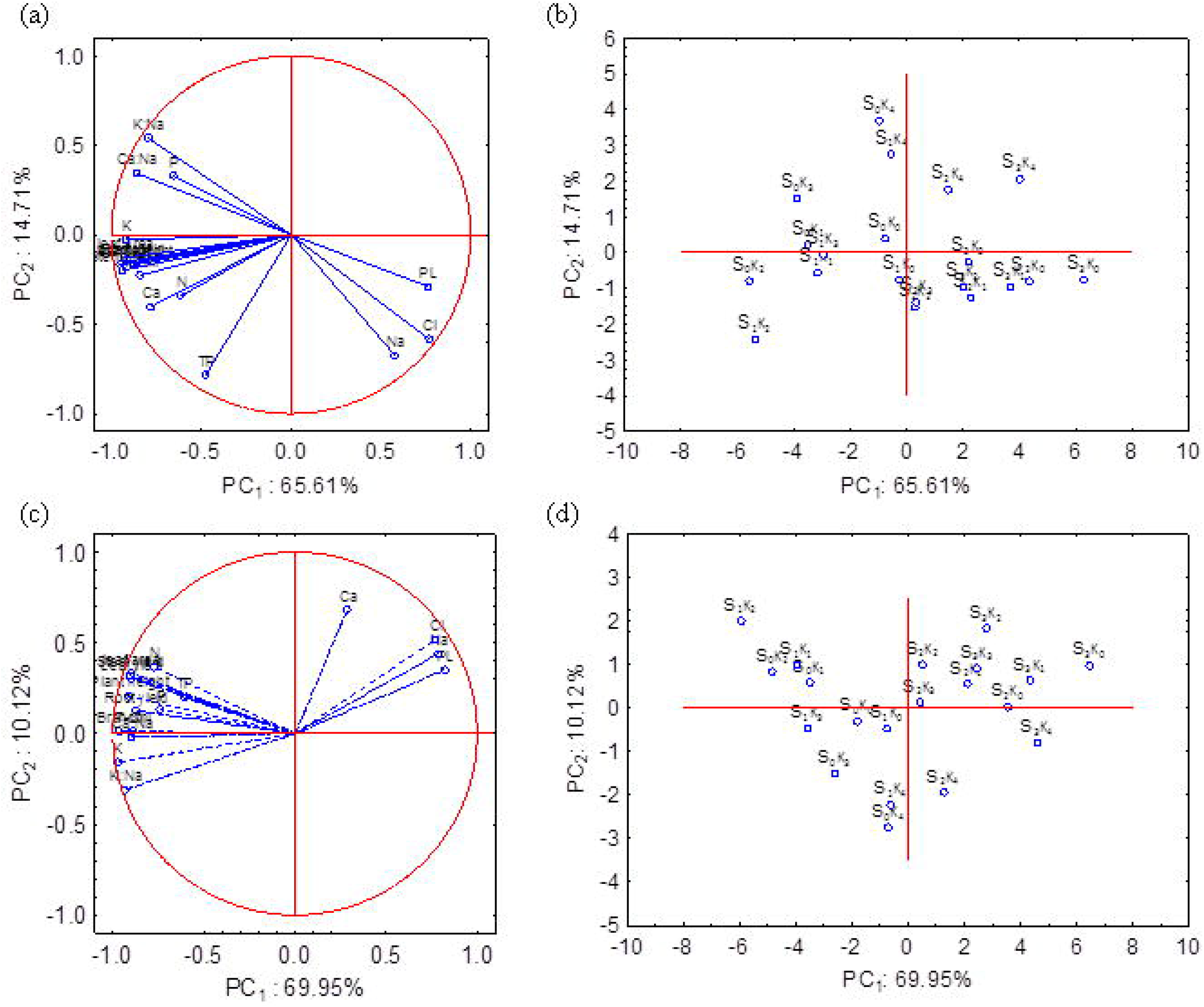
The multivariate analyses of agronomic traits (plant height, branch, leaf area, dry leaf, stem and root yield) and biochemical parameters [total chlorophyll (T-chl), total phenols (TP), proline (PL), N, P, K, Ca, Na, Cl, K: Na, Ca: Na] were conducted through Principal Component analysis (PCA). The first two factors (PC_1_ and PC_2_) mutually explained 80.32% and 80.07 % of the total variations for the 1^st^ and 2^nd^ year, respectively (**Fig. 6a-d**). The PCA bi-plots (**Fig. 6b and 6d**) represent the distributions of treatment combinations. The proline (PL), Na and Cl were placed in the positive coordinate of PC_1_ in 1^st^ year (**Fig. 6a**) with loading values of 0.82, 0.78 and 0.77, respectively. Similarly, in second cropping season PL, Na, Ca and Cl were also separated by the PC1 from rest of the agronomic traits and biochemical parameters, and placed in the positive coordinate of PC_1_ and PC_2_ (**Fig. 6c**). The PCA bi-plot (**Fig. 6b**) separated the treatments S_2_K_4_ (NaCl at 80 mM with the foliar application of KNO_3_ at 10.0 g L^−1^) and S_3_K_4_ (NaCl at 120 mM with foliar application of KNO_3_ at 10.0 g L^−1^) by PC_1_ and PC_2_, and placed in the positive coordinate of both PCs for 1^st^ season. However, no distinct group was formed in the PCA bi-plot (**Fig. 6d**) for 2^nd^ season.

Another PCA was carried out on steviol glycosides (stevioside, Reb-A, Reb-F, Reb-C, SB, TSGs) and biochemical traits (total Chl, total phenols, proline, N, P, K^+^, Ca^2+^, Na^+^, Cl^−^) to study the relationships among them. The PCA showed that the 1^st^ and 2^nd^ component (PC_1_ and PC_2_) cumulatively explained 61.62% and variations for 1^st^ and 2^nd^ cropping years, respectively (**Fig. 7 a-d**). The relationships among the variables (different steviol glycosides and biochemical traits) were presented in the space of the first two components (PC_1_ and PC_2_). During 1^st^ year Reb-F, Reb-C, proline, Na^+^ and Cl^−^ were separated from rest of variables by the PC_1_ and placed in negative coordinate (**Fig. 7a**). However, the PCA bi-plot (**Fig. 7b**) did not categorize the treatment combinations in any define clusters.

**Fig. 7.**
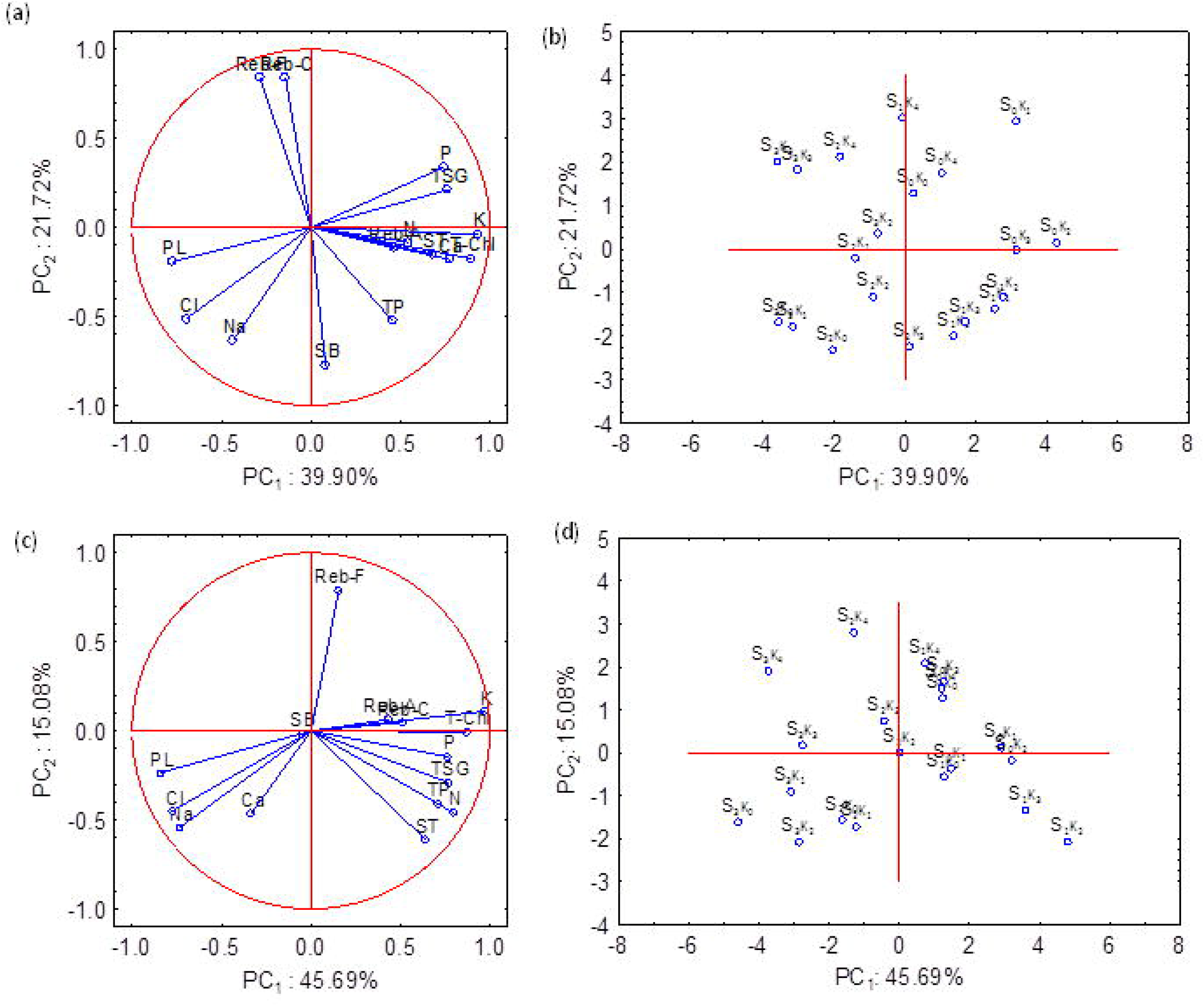
Principal Component analysis of secondary metabolites and biochemical traits. PC1 and PC_2_ jointly explains 61.62% and 60.77% of the total initial variability of the data in the 1^st^ (a, b) and 2^nd^ (c, d) cropping years, respectively (**Fig. 7 a-d**). During 1st year Reb-F, Reb-C, proline, Na+ and Cl-were separated from rest of variables by the PC1 and placed in negative coordinate (**Fig. 7a**). However, the PCA bi-plot (**Fig. 7b**) did not categorize the treatment combinations in any define clusters. Only S_0_K_1_ (irrigation with normal water X foliar application of KNO_3_ at 2.5 g L^−1^) was separated by PC_1_ and PC_2_, and situated in the positive coordinate of both PCs. Like 1st cropping season, Reb-F, Reb-C, proline, Na and Cl were again separated by the PC_1_ and placed in the negative coordinate (**Fig. 7c**) for 2^nd^ year. It was also revealed that the loading values three variables such as proline, Na and Cl were quite high with PC_1_. However, PCA bi-plot (**Fig. 7d**) separated the treatment combinations into two broad distinct clusters by PC_1_. Non-salinity (control) and low salinity (NaCl at 40 mM) treatments in combination with all foliar spray treatments (total 10 treatment combinations) were separated by PC_1_ and placed in positive coordinate (**Fig. 7d**). In this group, two treatment combinations namely S_1_K_2_ (NaCl at 40 mM X KNO_3_ at 5.0 g L^−1^) and S_1_K_3_ (NaCl at 40 mM × KNO_3_ at 7.5 g L^−1^) were further separated from rest of the treatment combinations by both PCs, and placed in positive coordinate of both PCs.

There were significant (*P* ≤ 0.05) differences in the soil pH and EC in response to different salinity levels in both the cropping seasons (**Fig. S1**). he status of available P, K^+^, Ca^2+^, Na^+^ and Cl^−1^ ions in the soil after harvest were significantly (P≤0.05) affected by the salinity levels in both the cropping seasons, whereas available N content was significantly (P≤0.05) affected only in 2^nd^ cropping season (**Table S2**).

## 4. Discussion

In salt-sensitive plant species, the growth inhibition is a common fact under saline conditions. In this experiment, plant height, number of branches and leaf area were significantly (*P* ≤ 0.05) reduced by NaCl at 120 mM. This result may be attributed to the accumulation of excess amount of Na^+^ in the older leaves, concurrently it reduces the uptake of other essential nutrients, such as potassium (K^+^), calcium (Ca^2+^), magnesium (Mg^2+^) and nitrate (NO_3_^−^) ions from the soil. Accumulation of excess amount of Na^+^ in the older leaves inhibits growth by accelerating their death consequently decreases the supply of carbohydrates to the meristematic regions (Munns 2002). In the present experiment, accumulation of Na^+^ in the leaves was increased by about 33-40% with NaCl at 120 mM compared with control (**Fig. 3a**).

The reduction in dry leaf yield with high salinity level (NaCl at 120 mM) was mainly due to the decrease of individual plant height, number of branch and LA per plant. Salinity decreases the ability of a plant to take up water by inducing osmotic stress and thereby reduces photosynthesis rate as a consequence of reduction of leaf expansion and closes stomates (Munns, 2005; Rahnama *et al.* 2010; Deinlein *et al.* 2014). Moreover, low biomass yield with high salinity level may be due to the fact that the high salinity induced oxidative stress, which damages membrane lipids, proteins and nucleic acids, and ultimately cell death (PérezLópez *et al.* 2009; Gill and Tuteja 2010; Del-Rio 2015). In our experiment, LA was reduced drastically at high salinity (NaCl at 120 mM) conditions probably due to the premature senescence of photosynthetic active leaves as a consequence of Na^+^ and Cl^−^ ions toxicity, which ultimately reduced the total dry leaf biomass. It has also been reported that under salt-stress a specific Cl^−^ build-up is observed in the leaves, which triggers 1-aminocyclopropane-1-carboxilic acid (ACC) synthesis and its conversion to ethylene, and eventually releases enough quantity of ethylene to hasten leaf abscission (Tudela and Primo-Millo 1992; Gómez-Cadenas *et al.* 1996, 1998; Dodd 2005).

The reduction of dry leaf biomass in the present experiment at high salinity stress (NaCl at 120 mM) can be due to the fact that the presence of higher concentration of NaCl in soils reduces the uptake of potassium (K^+^), calcium (Ca^2+^), and nitrogen (Fig 2a,e,g). Restrictions of N assimilation in plant body due to reduction of activity of cytosolic NADH nitrate reductase enzyme in nitrate assimilation pathway have been reported under salinity stress (Sivasankar and Oaks 1996; Jabeen and Ahmad 2011).

In the present study, the yield attributes and dry leaf biomass production of *S. Rebaudiana* plants did not show any reduction at mild salinity level (40 mM NaCl) rather to some extent increased compared with control (plants irrigated with non-saline water). These results could be due to the fact that Na^+^ played important role as a functional nutrient or manovalent cation in some physiological and metabolic activities and eventually hastened yield attributes of stevia. The functional role Na^+^ in the stomatal physiology of some plants has already been reported by Evans and Sorger (1966). Moreover, Cl^−^ accumulation in stevia plants at low salinity level (40 mM NaCl) may be the suitable amount for functioning in stomatal regulation and osmoregulation.

On the other hand, the rate of increase of dry leaf and stem yield with KNO_3_ at 5.0 g L^−1^ might be to the due to availability of adequate amount of K^+^ and its counter ion NO_3_^−^ for different biological activities like photosynthesis rate and RuBP carboxylase activity (Ramanujan and Rao 1971; Parasar and Dastane 1973; Jabeen and Ahmad 2011). Other probable explanation is higher accumulation of N in leaf, stem and root, which ultimately induces cytokinin synthesis in root tips and maintains desirable cytokinin and auxin ratio. The root cell division and differentiation of root are controlled by cytokinin and auxin ratio (Dello Ioio *et al.* 2008), and the events of major cell specification during embryogenesis are controlled by cytokinin and auxin (Muller and Sheen 2008). The positive effects of the N on yield attributes and yield of stevia plant have also been reported by Pal *et al.* (2013, 2015).

The dry leaf yield did not show significant reduction under moderate salinity (NaCl at 80 mM) when KNO_3_ at 10.0 g L^−1^ was applied compared with absolute control (plant irrigated with non-saline water and without KNO_3_). This result could be due to fact that exogenous application of K^+^ improved plant water status and maintained ion balance in cytosol, particularly K: Na and Ca: Na ratio. Higher K: Na ratio was observed with the exogenous application of K^+^ compared with water spray under all the salinity levels. Potassium play a key role in plant metabolic processes that improve plant water uptake by regulating the osmotic potential and hydraulic conductivity of membranes under drought or salinity stress (Heinen *et al.* 2009). Thus, it is evident from our investigation that exogenous application of K^+^ improves salinity tolerance of stevia upto 80 mM NaCl.

Reduction of photosynthetic pigments at higher salinity, such as Chl*a*, Chl*b* and total Chl in this experiment may be due to the accumulations of higher amount of Na^+^ and Cl^−^ in leaf, which disrupts the ultra structure of chloroplast and breaks down the Chl. Under salinity stress conditions, Chl degradation is happened due to enhanced activity of chlorophyllase enzyme (Santos 2004). However, some researcher has explained that Cl^−^ toxicity is the primary reason for the degradation of chlorophyll in plants (Tavakkoli *et al.* 2010). Chloroplasts also exhibit high permeability for Cl^−^ (Heber and Heldt 1981). On the other hand, the low ratio of Chl_*a*_: Chl_*b*_ with NaCl at 40 mM treatment in the present experiment provides evidence that the photosynthetic pigments are not degraded at mild salinity level. During the process of Chl degradation, Chl_*a*_ content is increased as a result of conversation from Chl_*b*_ to Chl_*a*_ (Fang *et al.* 1998; Eckardt 2009). Higher total Chl with KNO_3_ at 5.0 g L^−1^ could be due to the fact that right concentration of KNO_3_ increased the accumulation of total N and K^+^ in the leaf, and eventually increased the Chl content. K^+^ checks the decomposition of newly formed Chl and δ-aminolevulinic acid (ALA) formation (Tanaka and Tsuji 1980).

The organic osmolytes perform the vital role in maintaining low intracellular osmotic potential of plants and in preventing the detrimental effects of salinity stress (Tarczynski *et al.* 1993; Verslues *et al.* 2006). The utmost value of proline with NaCl at 120 mM could be due to fact that stevia plant faces the osmotic stress under salinity condition and proline is produced for osmotic adjustment. However, catabolism of proline is enhanced during recovery (Szekely *et al.* 2008; Poonlaphdecha *et al.* 2012; Sharma and Verslues 2010), and during this phase, proline regulates cell proliferation, cell death and expression of stress-recovery genes (Szabados and Savoure 2010). On the other hand, significantly (*P* ≤ 0.05) higher amount of proline was recorded with the foliar application KNO_3_ at 5.0 and 7.5 g L^−1^ compared with water spray in both the years.

It is well known fact that oxidative stress is occurred in salinity stress, mostly because of the generation of excessive ROS (reactive oxygen species) within plant cells. In order to scavenge or detoxify high ROS levels, plants produce phenolic compounds as an antioxidant defense system (Gill and Tuteja 2010; Foyer and Noctor 2011; Petridis *et al.* 2012; Chawla *et al.* 2013). Thus, biosynthesis of phenolic compounds is stimulated in salt-exposed plants (Navarro *et al.* 2006; Bose *et al.* 2014). However, in the present experiment, the total phenols content in stevia leaves was significantly increased (*P* ≤ 0.05) at mild salinity level (NaCl at 40 mM), whereas significant (*P* ≤ 0.05) reduction was observed at high salinity level (NaCl at 120 mM) compared with control (Fig. 2e). Maximum total phenols content with the foliar application of moderate concentration of KNO_3_ under low saline (NaCl at 40 mM) conditions (**Table 4**) may be due to accumulation of higher amount of K^+^ and balanced amount of others ions present in plant tissues, which are responsible for the accumulation of higher total phenols content. In the present experiment, higher amount of K^+^ in leaf tissues has been recorded with these treatment combinations. The growth strengthening antioxidant system in plants is improved by K^+^ under salinity stress (Zheng *et al.* 2008).

Averaged across the KNO_3_ levels, the plants treated with low salinity water (NaCl at 40 mM) accumulated higher amount of stevioside, Reb-A and TSGs compared with the plants treated with high salinity (NaCl at 120 mM) water and non-saline water in both the years. The present results are in conformity with the findings of Zeng *et al.* (2013) and Cantabella *et al.* (2017). Low SGs accumulation at severe salinity condition may be due to the fact that the energy is utilized for the process of maintaining metabolic homeostasis. It has been reported that when plants survive under high salinity stress conditions, energy is allocated for the synthesis of simple osmolytes and enhancing activities of antioxidant enzymes (Abrol *et al.* 2012). The role of SGs as osmoprotectant molecules under stress has already been established (Geuns and Ceunen 2013). On the other hand, irrespective of salinity levels, higher amounts of stevioside, Reb-A and TSGs in leaves were recorded with the moderate concentration of KNO_3_. This result may be due to the fact that exogenous application of KNO_3_ increases the accumulations of N, K^+^ and Ca^2+^ in plant tissues, which help to develop membrane system of chloroplasts and the content of photosynthetic pigments. Accumulation of steviol glycosides in the cells of stevia plant is correlated with the development of the membrane system of chloroplasts and the content of photosynthetic pigments (Ladygin *et al.* 2008). Moreover, under optimum nutrient availability, carbohydrates are used to increase the amount of SGs (Barbet-Massin *et al.* 2015).

Averaged over KNO_3_ level, concentration of ions particularly N, K^+^ and Ca^2+^ and ratio of K: Na and Ca: Na in plant tissues were drastically reduced at high salinity level (NaCl at 120 mM), whereas reversed trends were observed in case of Na^+^ and Cl^−^. In the saline soil, Na^+^ ion competes with K^+^ for the transporter, since both have the common transport mechanism (Sairam and Tyagi, 2004; Munns and Tester 2008) and similar in charge. However, at mild salinity (NaCl at 40 mM) level, K^+^ content in leaf was higher in plant tissue, which suggested that stevia could tolerate moderate salinity stress. This result also suggests that stevia plant has the ability to do selective K^+^ uptake under mild salinity level. Pooled across all salinity environments, the accumulations of N, K^+^ and Ca^2+^ in leaf, stem and root were higher with the foliar application of KNO_3_ at 5.0 gL^−1^. However, K^+^ accumulation was declined at higher concentration probably due to the destruction of ectodesmata structures (Marschner 1995). Moreover, K^+^ accumulation leaf tissue was increased with moderate concentration of KNO_3_ under all the salinity level. This may be due to fact that exogenous application of KNO_3_ increased the K^+^ accumulation to cope the adverse effect of salt stress.

Thus, it is concluded from the present investigation that the exogenous foliar application of KNO_3_ can elevate the salinity tolerance level of stevia through ion homeostasis, osmolyte accumulation, and antioxidant metabolism mechanism.

## Supplementary data

Table S1. Correlation coefficient matrix for agronomic traits of stevia

Table S2. Nutrients status in soil after harvest as influenced by salinity levels and foliar application of KNO_3_

Fig S1. Influence of irrigation water with different concentrations of NaCl on soil pH (a,b), EC (c,d) and organic carbon (e,f).

## Acknowledgments

The authors are grateful to Director of CSIR-IHBT, for providing the necessary facilities. We also thank Mr. Ramdeen Prasad for technical support. The authors acknowledge the CSIR, Government of India, for financial support.

## References

Abrol E, Vyas D, Koul S. 2012. Metabolic shift from secondary metabolite production to induction of antioxidative enzymes during NaCl stress in *Swertia chirata* Buch. -Ham. Acta Physiologia Plantarum 34, 541–546.

Arnon DI. 1949. Copper enzymes in isolated chloroplasts. Polyphenoloxidase in *Beta vulgaris*. Plant Physiology 24, 1–15.

Ashraf M, Foolad MR. 2007. Roles of glycine betaine and proline in improving plant abiotic stress resistance. Environmental and Experimental Botany 59, 206e216.

Barbet-Massin C, Giuliano S, Alletto L, Daydé J, Berger M. 2015. Nitrogen limitation alters biomass production but enhances steviol glycoside concentration in *Stevia rebaudiana* Bertoni. PLoS One 10(7), e0133067.

Bates LS, Waldren RP, Teare ID. 1973. Rapid determination of free proline for water stress studies. Plant soil 39, 205–207.

Bose J, Rodrigo-Moreno A, Shabala S. 2014. ROS homeostasis in halophytes in the context of salinity stress tolerance. Journal of Experimental Botany 65, 1241–1257.

Cantabella D, Piqueras A, Acosta-Motos JR, Bernal-Vicente A, Hernandez JA, Diaz-Vivancos P. 2017. Salt-tolerance mechanisms induced in *Stevia rebaudiana* Bertoni: Effects of mineral nutrition, antioxidative metabolism and steviol glycoside content. Plant Physiology and Biochemistry 115, 484–496.

Chawla S, Jain S, Jain V. 2013. Salinity induced oxidative stress and antioxidant system in salt-tolerant and salt-sensitive cultivars of rice *(Oryza sativa* L.). Journal of Plant Biochemistry and Biotechnology 22, 27–34.

Cornillon P, Palloix A. 1997. Influence of sodium chloride on the growth and mineral nutrition of pepper cultivars. Journal of Plant Nutrition 20, 1085–1094.

Deinlein U, Stephan AB, Horie T, Luo W, Xu G, Schroeder JI. 2014. Plant salt tolerance mechanisms. Trends in Plant Science 19, 371e379.

Del Rio LA. 2015. ROS and RNS in plant physiology: an overview. Journal of Experimental Botany 66(10), 2827–2837.

Dodd IC. 2005. Root-to-shoot signalling: Assessing the roles of ‘up’ in the up and down world of long-distance signalling in planta. Plant Soil 74, 257–275.

Eckardt NA. 2009. A new chlorophyll degradation pathway. Plant Cell 21, 700.

Evans HJ, Sorger G.J. 1966. Role of mineral elements with emphasis on the univalent cations. Annual Review of Plant Physiology 17, 47–76.

Fageria NK, Filho MPB, Moreira A, Guimares CM. 2009. Foliar fertilization of crop plants. Journal of Plant Nutrition 32, 1044–1064.

Fang Z, Bouwkamp J, Solomos T. 1998. Chlorophyllase activities and chlorophyll degradation during leaf senescence in nonyellowing mutant and wild type of *Phaseolus vulgaris* L. Journal of Experimental Botany 49, 503–510.

FAO. 2008. Land and Plant Nutrition Management Service. http://www.fao.org/ag/agl/agll/spush.

Foyer CH, Noctor G. 2011. Ascorbate and glutathione: the heart of the redox hub. Plant Physiology 155, 2–18.

Geuns JMC, Ceunen S. 2013. Steviol glycosides: chemical diversity, metabolism, and function. Journal of Natural Products 76, 1201e1228.

Gill SS, Tuteja N. 2010. Reactive oxygen species and antioxidant machinery in abiotic stress tolerance in crop plants. Plant Physiology and Biochemistry 48, 909–930.

Gómez-Cadenas A, Tadeo FR, Talon M, Primo-Millo E. 1996. Leaf abscission induced by ethylene in water stressed intact seedlings of Cleopatra mandarin requires previous abscisic acid accumulation in roots. Plant Physiology 112, 401–408.

Gómez-Cadenas A, Tadeo FR, Primo-Millo E, Talon M. 1998. Involvement of abscisic acid and ethylene in the response of citrus seedlings to salt shock. Physiologia Plantarum 103, 475–484.

Halperin ST, Gilroy S, Lynch JP. 2003. Sodium chloride reduces growth and cytosolic calcium, but does not affect cytosolic pH, in root hairs of *Arabidopsis thaliana* L. Journal of Experimental Botany 54, 1269–1280.

Heber U, Heldt HW. 1981. The chloroplast envelope: Structure, function, and role in leaf metabolism. Annual review of plant physiology 32, 132–168.

Heinen RB, Ye Q, Chaumont F. 2009. Role of aquaporins in leaf physiology. Journal of Experimental Botany 60(11), 2971e2985.

Husband AD, Godden W. 1927. Determination of sodium, potassium and chlorine in foodstuffs. Analyst 52, 72.

Dello Ioio R, Nakamura K, Moubayidin L, et al. 2008. A genetic framework for the control of cell division and differentiation in the root meristem. Science 332, 380–1384.

Izzo R, Navari-Izzo F, Quartacci MF. 1991. Growth and mineral absorption in maize seedlings as affected by increasing NaCl concentrations. Journal of Plant Nutrition 14, 687–699.

Jabeen N, Ahmad R. 2011. Foliar application of potassium nitrate affects the growth and nitrate reductase activity in sunflower and safflower leaves under salinity. Notulae Botanicae Horti Agrobotanici Cluj 39(2), 172–178.

James RA, Blake C, Byrt S, Munns R. 2011. Major genes for Na+ exclusion, Nax1 and Nax2 (wheat HKT1;4 and HKT1;5), decrease Na+ accumulation in bread wheat leaves under saline and waterlogged conditions. Journal of Experimental botany 62(8), 2939–2947.

Kinghorn AD. 2001. Stevia: The genus Stevia. pp. 1–202. Taylor & Francis Inc. New York, USA.

Ladygin VG, Bondarev NI, Semenova GA, Smolov AA, Reshetnyak OV, Nosov AM. 2008. Chloroplast ultrastructure, photosynthetic apparatus activities and production of steviol glycosides in *Stevia rebaudiana* in vivo and in vitro. Biologia Plantarum 52(1), 9–16.

Li L, Kim BG, Cheong YH, Pandey GK, Luan S. 2006. A Ca^2^+ signaling pathway regulates a K^+^ channel for low-K response in Arabidopsis. Proceedings of the National Academy of the Sciences, U. S. A 103, 12625–12630.

Marschner H. 1995. Mineral nutrition of higher plants. Annals of Botany 78, 527–528.

Muller B, Sheen J. 2008. Cytokinin and auxin interactions in root stem-cell specification during early embryogenesis. Nature 453, 1094–8.

Munns R, Tester M. 2008. Mechanisms of Salinity Tolerance. Annual Review of Plant Biology 59, 651–681.

Munns R. 2002. Comparative physiology of salt and water stress. Plant, Cell and Environment 25(2), 239–250.

Munns R. 2005. Genes and salt tolerance: bringing them together. New Phytologist 167(3), 645–63.

Navarro JM, Flores P, Garrido C, Martinez V. 2006. Changes in the contents of antioxidant compounds in pepper fruits at ripening stages, as affected by salinity. Food Chemistry 96, 66–73.

Niu X, Bressan RA, Hasegawa PM, Pardo JM. 1995. Ion homeostasis in NaCl stress environments. Plant Physiology 109(3), 735–742.

Pal PK, Mahajan M. 2017. Tillage system and organic mulch influence leaf biomass, steviol glycoside yield and soil health under sub-temperate conditions. Industrial Crops and Products 104, 33–44.

Pal PK, Kumar R, Guleria V, et al. 2015a. Crop-ecology and nutritional variability influence growth and secondary metabolites of *Stevia rebaudiana* Bertoni. BMC Plant Biology 15, 67.

Pal PK, Prasad R, Pathania V. 2013. Effect of decapitation and nutrient applications on shoot branching, yield, and accumulation of secondary metabolites in leaves of *Stevia rebaudiana* Bertoni. Journal of Plant Physiology 170, 1526–1535.

Parasar KS, Dastane NG. 1973. Studies on the effects of soil moisture regimes, plant population and nitrogen fertilization on the yield and quality of sugarbeet. Indian Journal of Agricultural Sciences 28, 516–520.

Perez-Alfocea F, Balibrea ME, Santa Cruz A, Estan MT. 1996 Agronomical and physiological characterization of salinity tolerance in a commercial tomato hybrid. Plant and soil 180, 251–257.

Perez-lopez U, Robredo A, Lacuesta M, Sgherri C, Muñoz-Rueda A, Navari-Izzo F, Mena Petite A. 2009. The oxidative stress caused by salinity in two barley cultivars is mitigated by elevated CO_2_. Physiologia Plantarum 135, 29–42.

Petridis A, Therios I, Samouris G, Tananaki C. 2012. Salinity-induced changes in phenolic compounds in leaves and roots of four olive cultivars *(Olea europaea* L.) and their relationship to antioxidant activity. Environmental and Experimental Botany 79, 37–43

Poonlaphdecha J, Maraval L, Roques S, Audebert A, Boulanger R, Bry X, Gunata Z. 2012. Effect of timing and duration of salt treatment during growth of a fragrant rice on variety on yield and 2-acetyl-1-pyrroline, proline, and GABA levels. Journal of Agricultural and Food Chemistry 60, 3824–3830.

Prasad R, Shivay YS, Kumar D, Sharma SN. 2006. Learning by doing exercise in soil fertility-a practical manual for soil fertility, Division of agronomy. New Delhi: IARI.

Rahnama A, James RA, Poustini K, Munns R. 2010. Stomatal conductance as a screen for osmotic stress tolerance in durum wheat growing in saline soil. Functional Plant Biology 37(3), 255–263.

Ramanujan T, Rao JS. 1971. Photosynthesis and dry matter production by rice plant growth under different levels of nitrogen. Madras Agricultural Journal 58, 38–40

Sairam RK, Tyagi A. 2004. Physiology and molecular biology of salinity stress tolerance in plants. Current Science 86(3), 407–421.

Santos CV. 2004. Regulation of chlorophyll biosynthesis and degradation by salt stress in sunflower leaves Among the amino acids, proline is the main effector in this response. Scientia Horticulturae 103, 93–99.

Schroeder JI, Delhaize E, Frommer WB, et al. 2013. Using membrane transporters to improve crops for sustainable food production. Nature 497, 60–66

Sharma S, Verslues PE. 2010. Mechanisms independent of abscisic acid (ABA) or proline feedback have a predominant role in transcriptional regulation of proline metabolism during low water potential and stress recovery. Plant Cell and Environment 33, 1838–1851.

Sivasankar S, Oaks A. 1996. Nitrate assimilation in higher plants: The effect of metabolites. Plant Physiology and Biochemistry 34, 609–620.

Subbarao GV, Johansen C, Jana MK, Kumar Rao JVDK. 1990. Effects of the sodium/calcium ratio in modifying salinity response of pigeon pea *(Cajanus cajan)*. Journal of plant physiology 136, 339–443.

Szabados L, Savoure A. 2010. Proline: a multifunctional amino acid. Trends in Plant Science 15, 89–97.

Szekely G, Abraham E, Cseplo A, et al. 2008. Duplicated *P5CS* genes of Arabidopsis play distinct roles in stress regulation and developmental control of proline biosynthesis. Plant Journal 53, 11–28.

Tanaka A, Tsuji H. 1980. Effects of cacium on chlorophyll synthesis and stability in the early phase of greening in cucumber cotyledons. Plant Physiology 65, 1211–5.

Tarczynski MC, Jensen RG, Bohnert HG. 1993. Stress protection of transgenic tobacco by production of the osmolyte mannitol. Science 259, 508–510.

Tavakkoli E, Fatehi F, Coventry S, Rengasamy P, McDonald GK. 2011. Additive effects of Na+ and Cl^−^ ions on barley growth under salinity stress. Journal of Experimental Botany 62(6), 2189–203.

Tavakkoli E, Rengasamy PSR, McDonald GK. 2010. High concentrations of Na+ and Cl^−^ ions in soil solution have simultaneous detrimental effects on growth of faba bean under salinity stress. Journal of Experimental Botany 61(15), 4449–4459.

Tudela D, Primo-Millo E. 1992. 1-Aminocyclopropane-1-carboxylic acid transported from roots to shoots promotes leaf abscission in *Cleopatra mandarin* (Citrus reshni Hort. ex Tan.) seedlings rehydrated after water stress. Plant Physiology 100, 131–137.

Verslues PE, Agarwal M, Katiyar-Agarwal S, Zhu J, Zhu JK. 2006. Methods and concepts in quantifying resistance to drought, salt and freezing, abiotic stresses that affect plant water status. Plant Journal 45, 523–539.

WHO. India. Available at: http://www.searo.who.int/india/mediacentre/events/2016/en/. Accessed on 4 My 2019.

Zeng J, Chen A, Dandan L, Bin Y, Wei W. 2013. Effects of salt stress on the growth, physiological responses, and glycoside contents of *Stevia rebaudiana* Bertoni. Journal of Agricultural and Food Chemistry 61, 5720–5726.

Zheng Y, Jia A, Ning T, Xu J, Li Z, Jiang G. 2008. Potassium nitrate application alleviates sodium chloride stress in winter wheat cultivars differing in salt tolerance. Journal of Plant Physiology 165(14), 1455e1465.

